# PeptiDIA: A Machine Learning Framework for Enhanced Peptide Identification in Fast-Gradient Data-Independent Acquisition Proteomics

**DOI:** 10.64898/2026.06.10.731224

**Authors:** Jordan Ortona, Mickaël Leclercq, Florence Roux-Dalvai, Bertrand Routy, Sébastien Bonnet, Arnaud Droit

## Abstract

Data-independent acquisition (DIA) mass spectrometry has become increasingly prevalent in proteomics as advances in instrumentation, chromatography, and computational analysis have enabled robust proteome identification across complex biological samples. However, analytical depth achieved with fast chromatographic gradients remains lower than that obtained using long-gradients, reflecting a throughput-depth trade-off. Here, we present PeptiDIA, a machine learning framework that enhances peptide identification in fast-gradient DIA data by leveraging paired fast and long-gradient acquisitions from identical samples. PeptiDIA processes DIA-NN outputs generated at relaxed false discovery rate thresholds to obtain expanded candidate peptide pools and trains gradient-boosted decision tree models using long-gradient identifications as reference labels. The model integrates DIA-NN features with engineered peptide descriptors and applies isotonic regression to calibrate probabilities, enabling controlled peptide recovery relative to the long-gradient reference. Applied to human and murine datasets spanning six tissues acquired on an Orbitrap Exploris 480, PeptiDIA increased peptide identifications by 25-34% at 1% target reference-discordance rate (RDR) and increased the number of protein groups containing at least one rescued peptide by 15-17%. Overall, PeptiDIA improves the identification depth of fast-gradient DIA-NN workflows without altering acquisition strategies. The framework is available as a web application and command-line tool at https://github.com/Jordano700/PeptiDIA.

## Introduction

Liquid chromatography coupled to tandem mass spectrometry (LC-MS/MS) is now the primary platform for large-scale peptide identification and protein quantification^1,2^. Over the past decade, data-independent acquisition (DIA) has largely replaced data-dependent acquisition (DDA) driven by the superior quantitative reproducibility and proteome coverage from DIA approaches^3–5^. In DIA, the mass spectrometer systematically fragments all precursor ions within predefined isolation windows across the entire mass range, generating comprehensive fragmentation spectra that are subsequently deconvolved using spectral libraries or library-free search algorithms^4^. This acquisition strategy eliminates stochastic sampling inherent to DDA methods and enables both deeper proteome coverage and more reproducible quantification across large sample cohorts^6^.

The analytical depth of DIA proteomics depends on chromatographic separation. In reversed-phase liquid chromatography, peptides are separated by hydrophobicity along an organic solvent gradient^7^, where gradient length determines peak capacity, the number of resolvable peaks within a separation window^8^. Longer gradients provide extended elution windows that reduce peptide co-elution, improve signal-to-noise ratios, and enable more confident peptide identification through better resolved chromatographic peaks^9^. Co-elution and chimeric spectra occur at all gradient lengths but worsen under compressed gradients, which also aggravate dynamic-range limitations as high-abundance signals mask co-eluting low-abundance peptides, complicating deconvolution and reducing identification confidence.

Standard DIA workflows employing 45-90 min LC gradients routinely achieve deep proteome coverage, identifying 6,000-8,000 protein groups from complex samples on platforms such as the Orbitrap Exploris 480^10,11^. However, this depth comes at limited throughput, expressed as samples per day (SPD). Long-gradient methods typically operate at 15-30 SPD, requiring 17-33 days of continuous instrument time to analyze a 500-sample cohort^12^. Fast-gradient DIA compresses separations to 5-15 min, increasing throughput to 100-300 SPD depending on configuration^13^. For example, the Evosep One system provides standardized methods ranging from 30 to 300 SPD, with the fastest completing cycles in under five minutes^14^. At 200 SPD, a 500-sample cohort can be processed in approximately 2.5 days. This acceleration reduces chromatographic resolution, increases peptide co-elution and spectral complexity, and typically lowers identification depth relative to matched long gradients^15^. Although the extent of reduction varies with sample complexity, instrument scan speed, and acquisition parameters, the throughput-depth trade-off remains a central unresolved challenge in DIA proteomics^4^.

Machine learning has become increasingly integrated into mass spectrometry-based proteomics^16,17,18^. Early approaches such as Percolator introduced semi-supervised learning for rescoring peptide-spectrum matches in DDA workflows, significantly improving identification sensitivity^19^. More recently, deep learning models have been developed to predict peptide fragmentation patterns, retention times, and collisional cross-sections, enabling enhanced library-free DIA analysis^20,21^. These approaches can integrate numerous variables simultaneously and recognize complex or subtle patterns that classical statistical approaches struggle to capture^22^.

Modern DIA analysis platforms now embed machine and deep learning directly within the search engine. DIA-NN, for example, employs neural network-based scoring and interference correction to enable deep, library-free peptide identification^23^. By implementing advanced statistical modeling and generating multiple quality metrics, DIA-NN improves identification confidence across diverse datasets^23^. However, its scoring framework is applied uniformly across chromatographic conditions, treating fast- and long-gradient acquisitions equivalently. This may be suboptimal for fast-gradient data, where peptide elution behavior and spectral characteristics differ systematically from longer separations, and even advanced search engines may reach their limits under compressed analytical conditions^24^.

Increasing throughput in DIA proteomics typically comes at the cost of identification sensitivity. We hypothesize that long-gradient acquisitions can provide high-confidence training labels to recover peptide identifications lost under fast-gradient conditions. By combining native DIA-NN features such as quality scores, spectral intensities, and chromatographic measurements with additional engineered features, supervised models can learn to recognize characteristic signatures of true peptides and recover identifications that fell below conventional confidence thresholds in fast-gradient searches.

Here, we present PeptiDIA, a machine learning framework that implements this paired-gradient learning strategy for enhanced peptide identification in fast-gradient DIA proteomics. PeptiDIA extracts features from DIA-NN search outputs and trains gradient-boosted decision tree models using long-gradient identifications as reference labels to enable calibrated peptide recovery at user-specified thresholds. We validate PeptiDIA using human and murine tissue datasets spanning six types, demonstrating consistent identification gains. The framework operates as a post-processing layer on DIA-NN search outputs and requires no modification to existing acquisition methods.

## Materials and Methods

### Overview of the PeptiDIA Framework

PeptiDIA is a supervised machine learning framework that rescores peptide candidates from fast-gradient DIA-NN searches using long-gradient identifications from the same samples as reference labels. The workflow comprises candidate generation at relaxed FDR thresholds, feature extraction and engineering, gradient-boosted decision tree classification, and probability calibration for controlled peptide recovery. Each stage is detailed in the sections below.

### Biological Samples and Experimental Design

The evaluation dataset comprised six tissue types from two species, providing biological diversity essential for robust model training and validation. Human tissues included artery (n=10), epicardial adipose (n=10), and heart (n=6, comprising 2 independent samples each analyzed in technical triplicates). Murine tissues included liver (n=10), ileum (n=10), and colon (n=10). Human biological samples were collected after obtaining written informed consent from all participants. The study protocol was reviewed and approved by the Research Ethics Committee of the Institut Universitaire de Cardiologie et de Pneumologie de Québec (IUCPQ) (CER20773, CER20841) and was conducted in accordance with applicable ethical guidelines and regulations. All animal experiments were approved by the Institutional Animal Care Committee of the Centre de Recherche du Centre Hospitalier de l’Université de Montréal (CRCHUM) and were conducted in accordance with the guidelines of the Canadian Council on Animal Care (C22055Br, C23028Br).

All samples represent independent biological specimens and were processed for tissue lysis, protein extraction, reduction, alkylation, and tryptic digestion according to three different protocols (Supplementary Methods).

### Mass Spectrometry analyses

Each biological sample was analyzed using two chromatographic regimes on an Orbitrap Exploris 480 mass spectrometer coupled to an Evosep One liquid chromatography system. Fast-gradient acquisitions employed the 300 SPD method, providing 4.8-minute gradients optimized for high-throughput. Long-gradient acquisitions used the 30 SPD method, delivering 48-minute gradients with enhanced chromatographic resolution. The mass spectrometer was operating in DIA mode with DIA window schemes and MS parameters optimized for each gradient length (Supplementary Methods).

### DIA-NN Data Processing

Raw mass spectrometry files were converted to mzML^25^ format using ThermoRawFileParser^26^ and subsequently processed with DIA-NN (version 2.20) in library-free mode with deep learning-based spectra prediction. Database searches were performed against UniProt reference proteomes (UP000005640 for human^27^; UP000000589 for mouse^28^). Search parameters included Trypsin/P digestion with one missed cleavage permitted, N-terminal methionine excision enabled, carbamidomethylation of cysteine as a fixed modification. Variable modifications were not included in the search (maximum variable modifications set to 0). Additional parameters included a peptide length range of 7-30 amino acids, precursor charge states 1-4, precursor m/z range of 300-1800, and fragment ion m/z range of 200-1800.

Each mzML file was processed as an independent DIA-NN analysis. Fast-gradient files were searched at 1%, 20%, and 50% FDR, whereas the paired long-gradient file from the same biological sample was searched at 1% FDR only and used as the high-confidence reference set. The 20% and 50% settings were retained intentionally as two permissive candidate-pool regimes (moderate and maximal expansion), enabling comparison of peptide recovery versus discordance-control behavior. The 50% threshold was used as the practical upper bound because thresholds above 50% did not yield additional candidates in our data.

DIA-NN reports results at the precursor level (one row per precursor ion, i.e., peptide at a specific charge state), so the same peptide can appear in multiple rows (e.g., +2 and +3). For peptide-level analysis, precursor rows were collapsed by Modified.Sequence (same modified peptide sequence across charge states), and precursor-level predictions were aggregated using the maximum prediction probability per Modified.Sequence. This max aggregation accepts a peptide if any of its precursor forms are confidently classified.

### Ground Truth Labeling Strategy

PeptiDIA defines peptide recovery as a binary classification task at the sample level. For each fast-gradient sample, candidate peptides from permissive searches (20% or 50% FDR) were labeled by comparison with the paired long-gradient 1% reference results: label = 1 if the Modified.Sequence was present, label = 0 if absent. Labels were charge-state invariant (presence at any charge state counted as present). Critically, peptides already identified at 1% FDR in the fast-gradient baseline were excluded from both training and testing, as these high confidence identifications require no recovery.

### Feature Selection and Engineering

For each precursor (peptide sequence + charge state) in the candidate pool, features were extracted across four categories capturing complementary aspects of identification confidence. DIA-NN output features were used directly from the parquet files generated by DIA-NN’s neural network scoring model. These included: (1) statistical quality metrics such as ‘Q.Value’ (precursor level), ‘Global.Q.Value’, posterior error probability (PEP), ‘PG.Q.Value’ (protein group), ‘GG.Q.Value’ (gene group), and evidence scores; (2) spectral measurements including MS1 precursor area, precursor m/z, charge state, apex intensity, best fragment m/z, and fragment m/z deviation; and (3) chromatographic metrics including observed and predicted retention times, peak width (FWHM), and elution profile characteristics, which are informative for fast-gradient data where co-elution affects identification confidence. PeptiDIA-engineered features were derived to augment the DIA-NN output. These included: (1) amino acid composition features comprising peptide length, counts of each of the 20 standard amino acids, and amino acid frequencies (count normalized by peptide length), encoding physicochemical properties that influence detectability; (2) log-transformed intensities using log1p to accommodate zero values and compress dynamic range; and (3) z-score normalizations for within-sample standardization of intensity-based features. The complete feature set comprised 87 features; a full list is provided in the Supplementary Materials (Table S1).

### Machine Learning Model

Gradient-boosted decision tree classifiers were implemented using XGBoost^29^. Hyperparameters were determined through grid search optimization across all tissue datasets (Table S2). For comparative model selection, alternative algorithms were evaluated including Random Forest, Logistic Regression, and Multi-Layer Perceptron (MLP) neural networks. To ensure fair comparison, hyperparameters for all models were independently optimized using Optuna, a Bayesian optimization framework, with 30 trials per model^30^. Model performance was assessed using Area Under the Receiver Operating Characteristic curve (AUC-ROC)^31^, which quantifies each classifier’s ability to discriminate between peptides validated by long-gradient ground truth (reference concordants) and peptides absent from the ground truth (reference discordants) across all possible classification thresholds. AUC-ROC was selected because it is threshold-independent and compares model performances without requiring probability-threshold calibration.

### Probability Calibration

Raw classifier scores were calibrated with isotonic regression^32^, which fits a monotonic mapping from model score to empirical class probability. Unlike parametric calibration (e.g., Platt scaling)^33^, isotonic regression makes no distributional assumption. Calibration used 3-fold stratified cross-validation on training data: out-of-fold predictions were generated for each fold, then pooled to fit the isotonic model. This avoids leakage and provides calibrated probabilities for downstream thresholding.

### RDR Threshold Determination

Because PeptiDIA is evaluated against paired long-gradient identifications rather than a target-decoy procedure, we report a **reference-discordance rate (RDR)** instead of conventional FDR.

At a probability threshold *t*, additional peptides are classified as reference-concordant (TP; present in the paired long-gradient 1% set) or reference-discordant (FP; absent from that set). RDR is defined as:

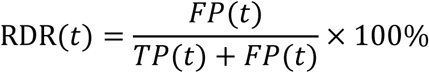

Target RDR values (e.g., 1%, 5%, 10%) are user-specified operating points, analogous to selecting an FDR threshold. For each target RDR, all unique calibrated probability thresholds were evaluated, and the threshold minimizing ∣ RDR(*t*) − Target_RDR ∣ was selected. Ties were resolved by retaining the threshold with higher peptide yield. Monotonic constraints were enforced so that higher target RDR values corresponded to more permissive thresholds and increased peptide counts. RDR was computed on additional peptides only, beyond the fast-gradient 1% baseline set.

We report results across a range of target RDR thresholds (1%, 2%, 5%, and 10%) rather than a single operating point because PeptiDIA is intended to expand overall peptide identification under fast-gradient conditions, and the appropriate level of stringency depends on the application. The 1% target RDR is the conservative default and is directly comparable to the 1% FDR convention in proteomics; it quantifies what can be recovered when discordance control is held stringent. The more permissive thresholds are intended for exploratory and discovery settings, such as screening a large cohort for candidate biomarkers, where maximizing peptide recovery is prioritized and missing a true signal is more costly than tolerating a higher proportion of reference-discordant identifications that can be confirmed in a subsequent targeted analysis. Reporting the full range characterizes the recovery-versus-discordance trade-off, supports a more robust assessment of the framework, and allows users to select an operating point matched to their study rather than imposing a single threshold.

### Validation Strategy

Two complementary validation protocols assessed model performance and generalizability, each evaluated using candidate pools generated at DIA-NN 20% and 50% FDR settings. For within-tissue validation (leave-one-sample-out), each biological sample was held out in turn as the test sample while the model was trained on all remaining biological samples from the same tissue. Splits were performed at the biological-sample level: for the heart dataset (two biological samples × three technical replicates each), all three technical replicates of a given biological sample were assigned together to either training or test, never split between them, to avoid leakage from shared biological background. Threshold calibration used 3-fold stratified cross-validation on training samples to generate out-of-fold predictions, from which RDR-calibrated thresholds were derived. For cross-tissue validation(leave-one-tissue-out), models were trained on all samples from five tissues and tested on all samples from the held-out sixth tissue, assessing generalization to unseen tissue types. Both validation strategies were conducted separately for FDR20 and FDR50 candidate pools, yielding four validation conditions: within-tissue FDR20, within-tissue FDR50, cross-tissue FDR20, and cross-tissue FDR50.

### Comparative Assessment of RDR against DIA-NN

To compare PeptiDIA with native DIA-NN scoring, we implemented a matched-yield analysis. DIA-NN assigns each precursor a q-value from target-decoy competition during the initial search, so raising the acceptance threshold beyond 1% is the most straightforward strategy for recovering additional peptides and the baseline we used for comparison. For each number of additional peptides recovered by PeptiDIA at a given RDR threshold, we identified the q-value cutoff producing the same yield and compared empirical RDR between methods.

### Quantification Validation

To confirm that recovered peptides exhibit biologically coherent quantitative behavior rather than representing noise, we assessed intensity correlations between gradient conditions. For each rescued peptide, precursor quantities (Precursor.Quantity from DIA-NN output) were extracted from both fast-gradient and paired long-gradient acquisitions. Intensities were log_2_-transformed prior to correlation analysis to normalize dynamic range. Spearman rank correlation coefficients (ρ) were calculated to measure monotonic relationships, as this metric is robust to intensity scale differences between gradient conditions^34^. Positive correlations indicate that rescued peptides maintain consistent quantitative relationships across acquisition conditions, supporting their biological validity independent of binary presence/absence classification. Furthermore, the ‘Quantity.Quality’ metric from DIA-NN output, which reflects the reliability of precursor quantification, was compared across all tissues and target RDR thresholds. Mann-Whitney U tests^35^ were used to assess statistical significance of quality score differences due to non-normal distribution.

### Feature Importance and Interpretability

Model interpretability was assessed through two complementary approaches. XGBoost’s built-in feature importance was calculated using information gain, which measures the average reduction in training loss contributed by each feature across all tree splits. Additionally, SHAP (SHapley Additive exPlanations) values were computed to quantify the marginal contribution of each feature to individual predictions^36^. Unlike gain-based importance, SHAP values provide directionality, indicating whether high or low feature values push predictions toward true positive or false positive classification and enable visualization of feature effects at the individual precursor level.

### Software Implementation and Availability

PeptiDIA was implemented in Python with core dependencies including XGBoost^29^, scikit-learn^37^, pandas^38^, and NumPy^39^. Two user interfaces are provided: a web application built on Streamlit^40^ enabling the complete workflow (dataset configuration, model training, inference, and result visualization) without programming requirements, and a command-line interface (CLI) that provides similar functionality within a terminal. Source code and documentation are available at https://github.com/Jordano700/PeptiDIA under the MIT open-source license.

### Computational Resources

All analyses were performed on a virtual machine equipped with an NVIDIA A100 GPU (40 GB VRAM). Leveraging GPU acceleration, PeptiDIA executes both model training and per-sample inference very rapidly (≈30 seconds to 1 minute). This enables the workflow to scale efficiently across large DIA cohorts, making it practical for high-throughput proteomics applications where hundreds of runs may be processed and reprocessed iteratively. In particular, the computational overhead of the machine learning step remains small relative to routine DIA data processing, allowing PeptiDIA to be integrated into analysis pipelines without becoming a bottleneck and supporting time-sensitive study designs that require fast turnaround.

**Figure 1.**
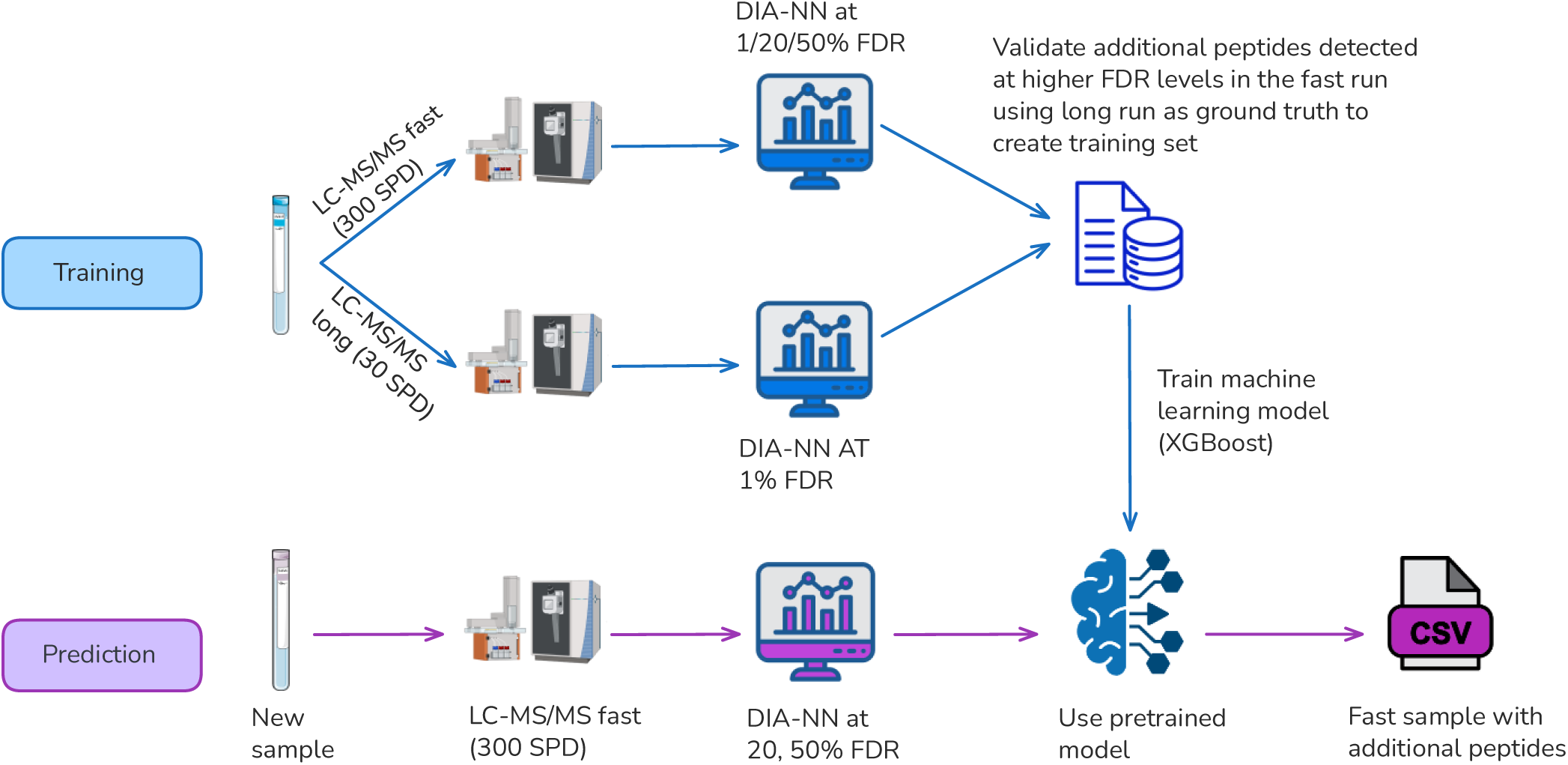
Overview of the PeptiDIA machine learning framework for enhanced peptide identification in high-throughput DIA proteomics. Samples are subjected to paired LC-MS/MS acquisitions using two chromatographic conditions: a fast-gradient (300 samples per day, SPD) optimized for high-throughput analysis, and a long-gradient (30 SPD) providing comprehensive peptide coverage. Data-independent acquisition (DIA) spectra are processed using DIA-NN. The fast-gradient is searched at multiple FDR thresholds (1%, 20%, and 50%), whereas the long-gradient is searched at 1% FDR establishing ground truth peptide identifications. This experimental design enables supervised learning: peptides detected at relaxed FDR thresholds in fast-gradient acquisitions are cross-referenced against the long-gradient ground truth to assign positive or negative labels. The resulting labeled dataset, comprising 87 features derived from DIA-NN output and feature engineering, is used to train an XGBoost gradient-boosted tree classifier with stratified 3-fold cross-validation. During inference, new fast-gradient samples are processed by DIA-NN at elevated FDR thresholds, and candidate peptides are rescored using the pre-trained model, enabling recovery of additional true identifications with controlled reference-discordant rates.

## Results

### Maximum Peptide Recovery Potential in Fast-Gradient DIA Data

We first established the ceiling for peptide recovery by comparing peptide identifications between paired fast-gradient (300 SPD) and long-gradient (30 SPD) acquisitions across 56 biological samples spanning six tissue types (Figure 2). The average number of identifications per tissue in the long-gradient runs are shown in Supplementary Figure S1.

**Figure 2.**
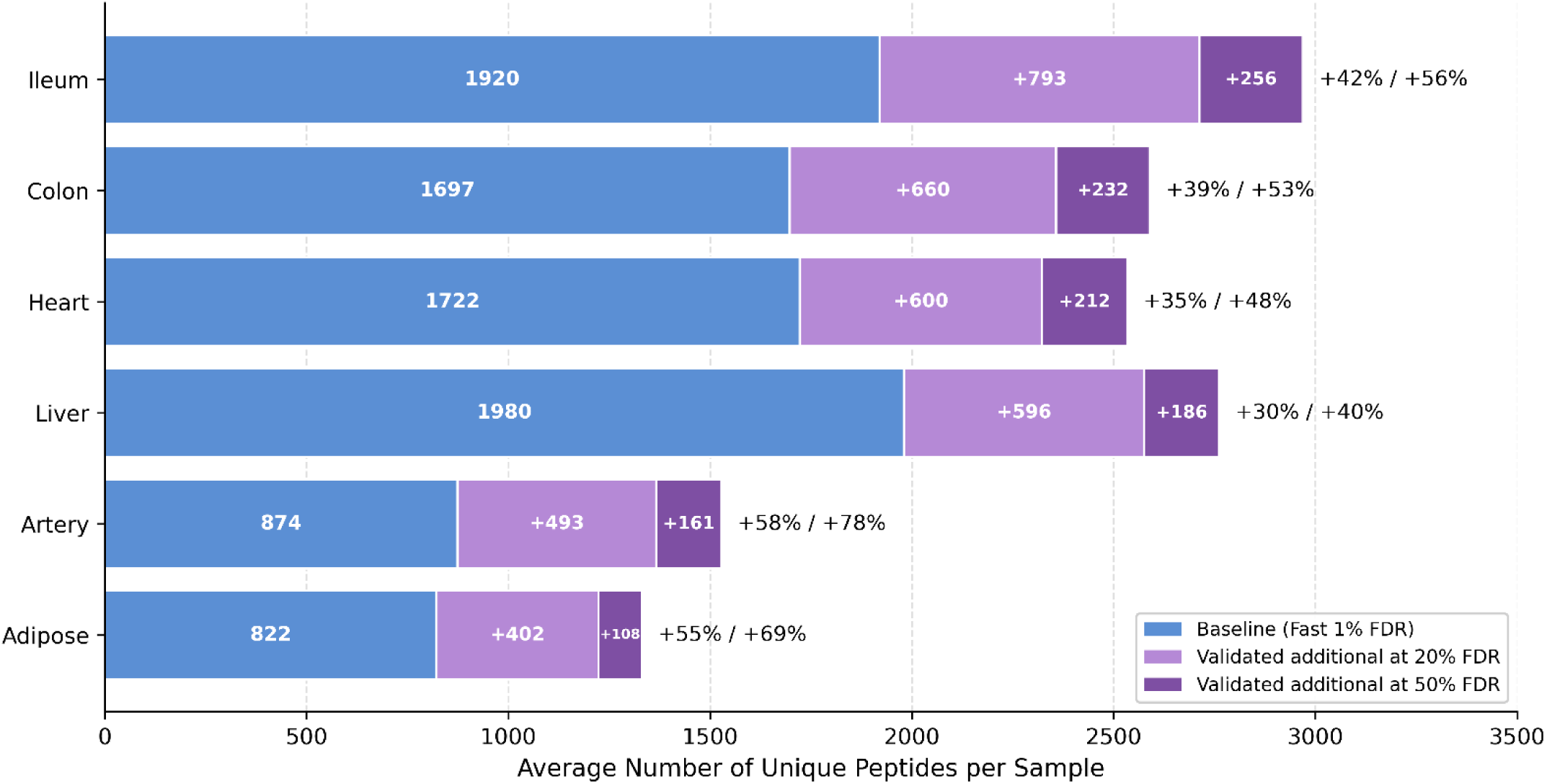
Maximum peptide recovery potential per tissue type. Stacked horizontal bars show the average number of unique peptides per sample across six tissue types (n = 6-10 samples per tissue). Blue segments represent baseline identifications from fast-gradient DIA (300 SPD, 1% FDR). Light purple segments indicate validated additional peptides detectable in fast-gradients at 20% FDR that are confirmed in the long-gradient ground truth (30 SPD, 1% FDR). Dark purple segments show further validated peptides when extending to 50% FDR. Percentage increases relative to baseline are shown for both thresholds (+20% FDR / +50% FDR). These validated candidates represent the maximum achievable through machine learning rescoring.

Baseline fast-gradient identifications at 1% FDR varied substantially across tissues, ranging from 822 ± 259 peptides per sample in adipose tissue to 1,980 ± 147 peptides in liver. Extending the DIA-NN search threshold from 1% to 20% FDR exposed additional validated candidates confirmed in the paired long-gradient ground truth, with a mean increase of +39.3% across tissues (range: +30.1% for liver to +58.0% for artery). Further extension to 50% FDR brought mean recoverable peptide increases to +52.1% (range: +39.5% for liver to +78.0% for artery).

These results establish that a perfect classifier operating on 20% or 50% FDR candidate pools could increase peptide identifications by around 40% or 60% on average across tissues respectively, relative to standard 1% FDR analysis. This substantial pool of validated peptides represents the maximum achievable gain through machine learning rescoring.

### PeptiDIA Increases Peptide Identifications Across Tissues and Training Configurations

Having established significant recovery potential in relaxed candidate pools, we evaluated PeptiDIA’s performance across within-tissue (leave-one-sample-out) and cross-tissue (leave-one-tissue-out) validation strategies using both FDR20 and FDR50 candidate pools (Figures 3-4, supplementary table S3). XGBoost consistently achieved superior AUC-ROC performance across all tissues and validation conditions and was therefore adopted as the primary classifier (Supplementary Figure S2).

**Figure 3.**
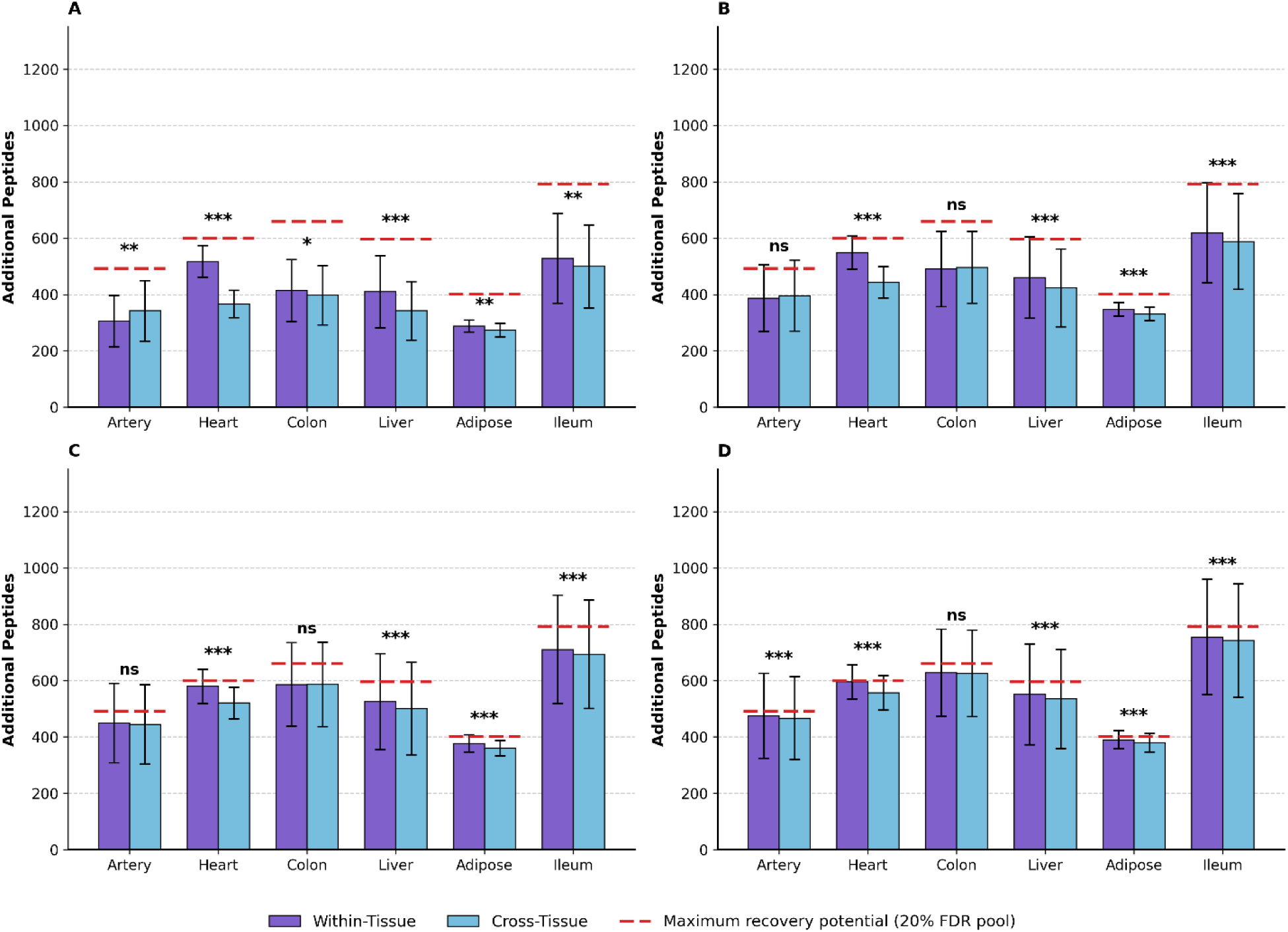
FDR20 model: Within-tissue and cross-tissue performance across target RDR thresholds. Bar plots showing the mean number of additional peptides recovered by PeptiDIA for each tissue at **(A)** 1%, **(B)** 2%, **(C)** 5%, and **(D)** 10% target RDR. Machine learning models were trained using additional peptides recovered by running DIA-NN at 20% FDR. Within-tissue models (purple) were trained on n−1 samples from the same tissue and tested on the held-out sample. Cross-tissue models (blue) were trained on all samples from the 5 other tissues and tested on the held-out tissue. Red dashed lines mark the maximum recovery potential for each tissue, defined as the mean number of validated additional peptides available in the 20% FDR candidate pool (the per-tissue ceiling shown in Figure 2). Error bars represent standard deviation. Significance markers indicate paired t-test results (within-tissue vs. cross-tissue, paired by sample): ns = not significant, *p < 0.05, **p < 0.01, ***p < 0.001.

**Figure 4.**
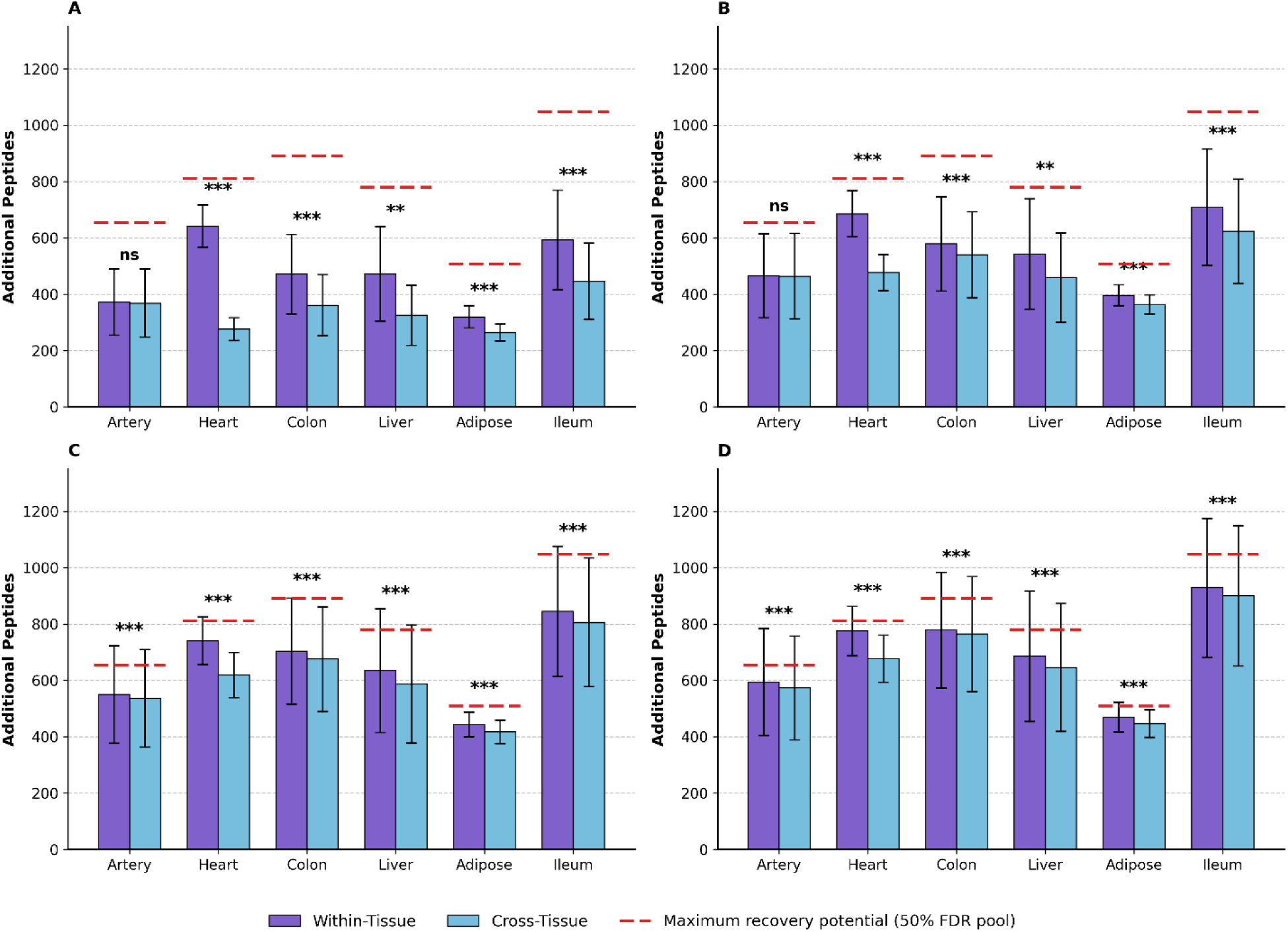
FDR50 model: Within-tissue and cross-tissue performance across target RDR thresholds. Bar plots showing the mean number of additional peptides recovered by PeptiDIA for each tissue at **(A)** 1%, **(B)** 2%, **(C)** 5%, and **(D)** 10% target RDR. Machine learning models were trained using additional peptides recovered by running DIA-NN at 50% FDR. Within-tissue models (purple) were trained on n−1 samples from the same tissue and tested on the held-out sample. Cross-tissue models (blue) were trained on all samples from the 5 other tissues and tested on the held-out tissue. Red dashed lines mark the maximum recovery potential for each tissue, defined as the mean number of validated additional peptides available in the 50% FDR candidate pool (the per-tissue ceiling shown in Figure 2). Error bars represent standard deviation. Significance markers indicate paired t-test results (within-tissue vs. cross-tissue, paired by sample): ns = not significant, *p < 0.05, **p < 0.01, ***p < 0.001.

#### Within-Tissue Validation

At 1% target RDR, within-tissue models achieved mean peptide increases of +30.0% (FDR20 training) and +33.6% (FDR50 training) across tissues. Recovery ranged from adipose (lowest: 288-320 additional peptides) to ileum/heart (highest: 528-642 peptides), with FDR50 training consistently outperforming FDR20 by 12-25% depending on tissue. Heart showed the largest improvement from expanded training pools (642 vs 517 peptides, +24.2%). At 10% target RDR, maximum mean recovery reached +41.8% (FDR20) to +49.7% (FDR50) additional peptides.

#### Cross-Tissue Validation

Cross-tissue models, which assess generalization to tissue types absent from training, achieved slightly lower but substantial gains. At 1% target RDR, cross-tissue models achieved +25.2-27.9% mean peptide increases (FDR50-FDR20), representing 75-85% of within-tissue performance. Performance scaled with target RDR threshold, reaching +40.7-47.3% at 10% target RDR. Within-tissue models significantly outperformed cross-tissue models for most tissues at 1% target RDR (paired t-test, p < 0.01-0.001), with artery typically showing no significant difference between validation strategies; at 10% target RDR, within-tissue advantages reached significance for all six tissues. The red bars in Figures 3 and 4 mark the per-tissue recovery ceiling from Figure 2. At 1% target RDR, the operating point most relevant for routine proteomics, within-tissue models attained a mean of 69% (FDR20) and 60% (FDR50) of the recoverable pool, while cross-tissue models, which never see the target tissue during training, attained 63% (FDR20) and 43% (FDR50). Recovered peptides converged toward the ceiling as the threshold relaxed: at 10% target RDR, within-tissue models reached 96% (FDR20) and 90% (FDR50), and cross-tissue models reached 93% (FDR20) and 85% (FDR50).

#### Summary of Recovery Performance

Table 1 summarizes mean percentage increases across all configurations (per-tissue breakdowns in Table S3). At 1% target RDR, peptide increases ranged from 25.2-33.6% while corresponding protein groups with at least one rescued peptide increased by 15.4-17.1%. Protein groups are derived from DIA-NN Protein.Group annotations and are not reported with protein-level FDR control. These gains were consistent across higher RDR thresholds, reaching maximum peptide increases of 41.8-49.7% at 10% target RDR.

**Table 1.** Mean Peptide and Protein Group Percentage Increases Across Training Strategies. Mean percentage increase across 6 tissues compared to DIA-NN baseline (1% FDR, fast-gradient search). Peptide increase: (additional peptides / baseline peptides) × 100. Protein group with at least one rescued peptide increase: novel protein groups identified from DIA-NN Protein.Group annotations, calculated as (new protein groups / baseline protein groups) × 100. **(A, C) Cross-tissue:** Models trained on 5 other tissues. **(B, D) Within-tissue:** Models trained on n−1 samples from same tissue. **FDR20/FDR50:** FDR selected on DIA-NN (20% or 50%) used to recover additional peptides; both training and inference datasets contain peptides at this threshold.

| (A) FDR20 Cross-Tissue |  |  | (B) FDR20 Within-Tissue |  |  |
| --- | --- | --- | --- | --- | --- |
| Target RDR | Peptide Increase (%) | Protein Increase (%) | Target RDR | Peptide Increase (%) | Protein Increase (%) |
| 1% | 27.9 | 16.4 | 1% | 30.0 | 16.8 |
| 2% | 33.5 | 17.0 | 2% | 35.3 | 17.2 |
| 5% | 38.4 | 17.3 | 5% | 39.8 | 17.4 |
| 10% | 40.7 | 17.4 | 10% | 41.8 | 17.5 |

**Table 1. Mean Peptide and Protein Group Percentage Increases Across Training Strategies.**
| (C) FDR50 Cross-Tissue |  |  | (D) FDR50 Within-Tissue |  |  |
| --- | --- | --- | --- | --- | --- |
| Target RDR | Peptide Increase (%) | Protein Increase (%) | Target RDR | Peptide Increase (%) | Protein Increase (%) |
| 1% | 25.2 | 15.4 | 1% | 33.6 | 17.1 |
| 2% | 35.5 | 17.1 | 2% | 39.9 | 17.5 |
| 5% | 43.3 | 17.7 | 5% | 46.2 | 17.8 |
| 10% | 47.3 | 17.8 | 10% | 49.7 | 18.0 |

### RDR Calibration Enables Reliable Discordance Control on Recovered Peptides

The practical utility of PeptiDIA depends on reliable control over discordant identifications among recovered peptides. We evaluated calibration accuracy by comparing empirical RDR against target operating points (Supplementary Figures S3-S4). At stringent thresholds (1-2% target RDR), calibration was accurate across all model configurations. FDR20 models achieved mean actual RDR of 0.97-1.08% at 1% target and 2.17-2.40% at 2% target. FDR50 models showed slightly conservative calibration at lower thresholds (0.52-0.55% actual RDR at 1% target), ensuring users do not exceed their specified reference discordance tolerance. At higher thresholds (5-10%), calibration showed increased deviation from target, particularly for within-tissue models where actual RDR ranged from 5.7-8.9% at 5% target and 12.0-18.6% at 10% target. Users requiring stringent RDR control should apply conservative thresholds (1-2%)

### PeptiDIA Achieves better RDR Control Compared to DIA-NN Q-Value Relaxation

To compare PeptiDIA against native DIA-NN scoring, we performed matched-yield comparisons across all validation conditions (Table 2; per-tissue breakdowns provided in Table S5). At 1% target RDR, PeptiDIA achieved substantially lower empirical RDR than DIA-NN q-value thresholding across all model configurations. Within-tissue models recovered 411-479 additional peptides (FDR20-FDR50) with 1.1% actual RDR, compared to 17.4-19.9% for matched-yield q-value relaxation. Cross-tissue models recovered 340-371 additional peptides with 1.0-1.3% actual RDR, compared to 15.2-16.1% for q-value relaxation.

**Table 2.** Mean ML RDR vs DIA-NN FDR Across All Tissues. Mean additional peptides recovered (Add. Pep.) and actual RDR achieved by PeptiDIA (ML RDR) at each target RDR threshold, averaged across six tissues. To enable direct comparison, the DIA-NN Q-value threshold was increased until the same number of additional peptides was recovered, and the resulting RDR is reported (DIA-NN FDR). **FDR20/FDR50:** DIA-NN FDR (20% or 50%) used to define the candidate peptide pool for machine learning classification. **Cross-Tissue:** Models trained on samples from five tissues and applied to the held-out sixth tissue. **Within-Tissue:** Models trained on n-1 samples from the same tissue and applied to the remaining held-out sample.

| Model | Target RDR | Add. Pep. | ML RDR (%) | DIA-NN FDR (%) |
| --- | --- | --- | --- | --- |
| **FDR20 Cross-Tissue** | 1% | 371 | 1.3 | 16.1 |
|  | 2% | 447 | 2.2 | 18.9 |
|  | 5% | 518 | 5.3 | 21.6 |
|  | 10% | 551 | 10.3 | 22.8 |
| **FDR20 Within-Tissue** | 1% | 411 | 1.1 | 17.4 |
|  | 2% | 476 | 2.6 | 19.9 |
|  | 5% | 538 | 7.1 | 22.4 |
|  | 10% | 566 | 14.6 | 23.4 |
| **FDR50 Cross-Tissue** | 1% | 340 | 1.0 | 15.2 |
|  | 2% | 488 | 1.9 | 20.4 |
|  | 5% | 607 | 5.0 | 25.2 |
|  | 10% | 668 | 9.7 | 27.8 |
| **FDR50 Within-Tissue** | 1% | 479 | 1.1 | 19.9 |
|  | 2% | 563 | 2.4 | 23.3 |
|  | 5% | 653 | 6.5 | 27.2 |
|  | 10% | 706 | 13.5 | 29.5 |

This pattern was consistent across all target RDR levels. At 2% target RDR, PeptiDIA achieved 1.9-2.6% actual RDR compared to 18.9-23.3% for q-value relaxation. At 5% target RDR, PeptiDIA achieved 5.0-7.1% actual RDR compared to 21.6-27.2%. At 10% target RDR, PeptiDIA achieved 9.7-14.6% actual RDR compared to 22.8-29.5%. Across all conditions, PeptiDIA reduced empirical RDR by 10-18 percentage points relative to q-value relaxation at equivalent peptide yield.

### Recovered Peptides Exhibit Quantitative Coherence with Ground Truth

To confirm that recovered peptides represent genuine biological signal, we assessed quantitative coherence through cross-gradient intensity correlations and DIA-NN’s Quantity.Quality metric.

#### Cross-Gradient Intensity Correlations

PeptiDIA-recovered peptides exhibited strong quantitative agreement between fast-gradient and paired long-gradient acquisitions (Table 3; per-tissue correlations in Table S6). Baseline peptides (identified at 1% FDR in the fast-gradient) showed mean correlation of ρ = 0.846 across tissues. At 1% target RDR, recovered peptides maintained moderate correlations with ground truth, below the baseline value. FDR20 within-tissue models achieved mean ρ = 0.640, FDR20 cross-tissue ρ = 0.653, FDR50 within-tissue ρ = 0.626, and FDR50 cross-tissue ρ = 0.660. At 2% target RDR, correlations decreased modestly: FDR20 within-tissue ρ = 0.580, FDR20 cross-tissue ρ = 0.600. At 5% target RDR, FDR20 within-tissue achieved ρ = 0.466, FDR20 cross-tissue ρ = 0.531. At 10% target RDR, correlations decreased further: FDR20 within-tissue ρ = 0.357, FDR20 cross-tissue ρ = 0.377.

**Table 3.** Mean Quantification Validation Across All Tissues. Mean Spearman correlation coefficients (ρ) between log₂-transformed precursor intensities of peptides identified in fast-gradient (300 SPD) runs and matched ground truth intensities from long-gradient (30 SPD) runs, averaged across six tissues). **Baseline:** High-confidence peptides identified at 1% FDR in the fast-gradient. **1-10% RDR:** ML-rescued peptides at each target RDR threshold. **FDR20/FDR50:** DIA-NN FDR (20% or 50%) used to define the candidate peptide pool for machine learning classification. **Within-Tissue:** Models trained on n-1 samples from the same tissue and tested on the held-out sample. **Cross-Tissue:** Models trained on all samples from 5 other tissues and tested on the held-out tissue.

| Model | Baseline | 1% RDR | 2% RDR | 5% RDR | 10% RDR |
| --- | --- | --- | --- | --- | --- |
| **FDR20 Within-Tissue** | 0.846 | 0.640 | 0.580 | 0.466 | 0.357 |
| **FDR20 Cross-Tissue** | 0.846 | 0.653 | 0.600 | 0.531 | 0.377 |
| **FDR50 Within-Tissue** | 0.846 | 0.626 | 0.514 | 0.327 | 0.246 |
| **FDR50 Cross-Tissue** | 0.846 | 0.660 | 0.597 | 0.431 | 0.281 |

#### Quantity.Quality Score Distributions

To further evaluate the quantification quality of peptides recovered by PeptiDIA, we examined DIA-NN’s Quantity.Quality metric. Supplementary Figures S5-S6 present these distributions across tissues and model configurations.

Baseline peptides (identified at 1% FDR in fast-gradient) exhibited the highest mean Quantity.Quality scores (∼0.75-0.85 across tissues), reflecting the stringent confidence threshold applied during initial identification. True positive peptides recovered by PeptiDIA showed intermediate quality scores (∼0.50-0.60), while false positive peptides showed systematically lower scores (∼0.40-0.45). This hierarchy was consistent across all tissues and model configurations (Mann-Whitney U test, p < 0.001 for all TP vs FP comparisons).

### Feature Importance Reveals Discriminative Variables

XGBoost gradient-boosted decision tree models enable calculation of feature importance through information gain contributions. Analysis of feature importance was performed across all trained models for both FDR20 and FDR50 candidate pools under within-tissue and cross-tissue conditions (Figure 5; full feature importance rankings provided in table S7). DIA-NN protein-level quality metrics consistently dominated feature importance rankings across all conditions. Global.Protein.Q.Value was the most discriminative feature, followed closely by Protein.Q.Value and Protein.PEP, which ranked among the top four features in all models. Gene.Group.Q.Value also contributed substantially, particularly in FDR50 models. Amino acid composition features provided secondary discriminative power. Cysteine count and cysteine frequency were the top-ranked amino acid features across all conditions, followed by methionine count and methionine frequency. Other contributing features included Precursor.Charge, Proteotypic status, and Global.Peptidoform.Q.Value.

**Figure 5.**
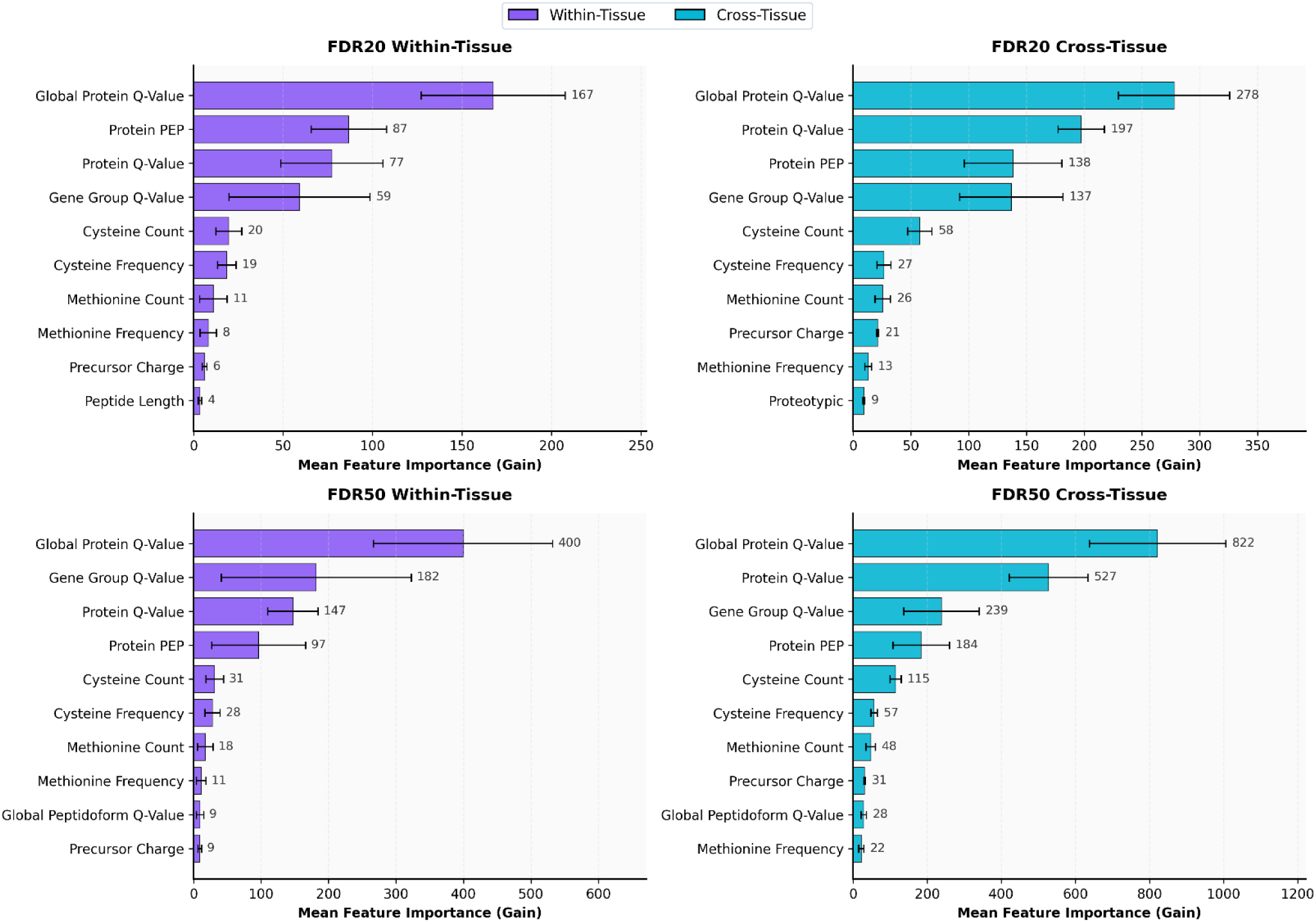
Feature importance analysis of XGBoost models across validation conditions. Mean feature importance (gain) for the top contributing features in FDR20 Within-Tissue (top left), FDR20 Cross-Tissue (top right), FDR50 Within-Tissue (bottom left), and FDR50 Cross-Tissue (bottom right) models. Gain represents the average reduction in training loss contributed by a feature across all tree splits.

#### SHAP Analysis

SHAP analysis revealed the directionality of feature contributions at the individual peptide level (Supplementary Figure S7). Lower protein-level q-values consistently pushed predictions toward true positive classification, while higher q-values pushed toward false positive classification. Among amino acid composition features, lower cysteine content and shorter peptide lengths were associated with true positive predictions. These directional patterns were consistent across all four model configurations (FDR20/FDR50, within-tissue/cross-tissue).

## Discussion

### Unlocking Hidden Identification Potential in Fast-Gradient Data

Our initial analysis revealed that fast-gradient DIA spectra contain substantially more genuine peptide signals than conventional 1% FDR filtering recovers. When we extended DIA-NN searches to relaxed FDR thresholds and validated candidates against paired long-gradient ground truth, we found that recoverable peptides exceeded 50% above baseline for several tissues. These results suggest that stringent FDR thresholds sacrifice true identifications that possess genuine spectral evidence but fall below confidence cutoffs due to spectral complexity arising from chromatographic compression. Benchmarking against this recovery ceiling shows that even cross-tissue models, which never see the target tissue during training, recover the majority of the theoretically attainable peptides at the stringent 1% target RDR used in routine proteomics, and approach the ceiling almost completely as the threshold is relaxed; within-tissue models recover an even larger share. PeptiDIA therefore captures most of the genuine signal hidden in fast-gradient data rather than leaving it unidentified.

### Consistent Recovery Across Diverse Biological Contexts

The consistent performance across six tissue types from two species demonstrates PeptiDIA’s robustness to biological heterogeneity. Adipose, artery, heart, colon, ileum, and liver represent distinct proteome compositions, dynamic ranges, and analytical challenges, yet the framework achieved significant gains in each context. The greater recovery achieved by FDR50 models compared to FDR20 models across all tissues indicates that widening candidate pools provides richer training signal. We interpret this finding as evidence that exposing the classifier to peptides across the full spectrum of spectral quality, from confidently identified to marginally detectable enables learning of subtle discriminative signatures that separate genuine identifications from interference. Laboratories should process fast-gradient data at maximally permissive thresholds when generating candidate pools for PeptiDIA rescoring.

### Cross-Tissue Generalization Enables Broad Deployment

The modest performance gap between within-tissue and cross-tissue models suggests that the features driving classification, particularly DIA-NN quality metrics and amino acid composition, capture fundamental biochemical properties of peptide detectability that transcend tissue-specific proteome composition. Laboratories can therefore use existing multi-tissue models rather than generating tissue-specific training sets, which is particularly relevant for workflows processing diverse sample types.

### Rigorous RDR Control Distinguishes Machine Learning from Threshold Relaxation

The matched-yield comparison demonstrates that PeptiDIA achieves markedly lower empirical RDR than q-value relaxation at equivalent peptide yield across all conditions. This difference arises because raising the q-value threshold indiscriminately accepts all precursors above the cutoff, whereas PeptiDIA evaluates each candidate using multiple orthogonal features, selectively recovering peptides with genuine spectral evidence while rejecting co-scoring false positives. Isotonic regression calibration further ensures actual discordance rates closely match user-specified RDR targets.

### Quantitative Validity Supports Downstream Analysis

Identification gains are only meaningful if recovered peptides support accurate quantification. Intensity correlations for recovered peptides were moderate and lower than for baseline identifications, a difference that is largely explained by abundance: the peptides PeptiDIA recovers are predominantly lower-abundance species, approximately 2-fold lower than baseline identifications on average at 1% RDR, with this difference increasing at higher target RDR thresholds (Supplementary Figure S8), whose weaker signals are more susceptible to run-to-run variability. This decline in correlation at higher target RDR thresholds reflects a sensitivity/precision trade-off that users can calibrate to their application. Together with the enrichment of higher ‘Quantity.Quality’ scores among true positives, this suggests that the classifier recovers peptides with reliable quantitative signal rather than noise. As signal intensity decreases, the relative contribution of instrumental noise increases, reducing measurement reproducibility⁴¹. The observed correlation decrease therefore reflects the abundance distribution of rescued peptides rather than errors in identification.

### Interpretable Features Enable Biological Insight

DIA-NN quality metrics dominate feature importance rankings, indicating that PeptiDIA effectively leverages existing search engine confidence signals. Critically, these metrics, including columns labeled ‘Global’, were computed within each single-file DIA-NN analysis and were not derived from a multi-run cohort-wide search. This distinction matters because cohort-level processing would alter the distribution of confidence scores across files, potentially inflating the apparent importance of these features. The most discriminative feature was ‘Global.PG.Q.Value’, a protein-group-level false discovery rate estimate. DIA-NN computes two tiers of q-values using distinct statistical procedures²³: run-specific q-values (’Q.Value’, ‘PG.Q.Value’) derived from within-run target-decoy competition, and global q-values (’Global.Q.Value’, ‘Global.PG.Q.Value’) designed for experiment-wide confidence but computed using a separate scoring algorithm that also produces distinct values when applied to single runs²³. In our data, ‘Global.Q.Value’ exceeded ‘Q.Value’ for every precursor while preserving identical rank order (Spearman ρ = 1.0). The greater feature importance of global q-values likely reflects their wider dynamic range among candidate peptides: run-specific q-values cluster near zero for precursors that passed or nearly passed the 1% FDR threshold, whereas the more conservative global estimates spread these same candidates across a broader score range, providing the classifier with greater discriminative power for tree-based splitting. Both tiers appear among top-ranked features, alongside gene group q-values (’GG.Q.Value’) and posterior error probability (’PEP’), indicating that PeptiDIA learns to exploit discrepancies between these metrics. A peptide with a favorable precursor q-value but unfavorable protein group q-value, or one where global and run-specific estimates diverge, carries identification reliability information that no single threshold captures.

The secondary contribution of amino acid composition, particularly cysteine and methionine, aligns with known biochemistry. These residues influence ionization efficiency, fragmentation patterns, and chromatographic behavior in ways that affect identification confidence under fast-gradient conditions⁴². Cysteine’s fixed carbamidomethyl modification, which alters its mass, ionization, and fragmentation (and is occasionally incomplete, leaving some cysteines uncapped), and methionine’s susceptibility to oxidation exhibit distinctive behavior that may amplify identification challenges when chromatographic separation is compressed⁴³. The interpretability of these features distinguishes PeptiDIA from black-box approaches and supports that the classifier captures meaningful biochemical patterns rather than dataset-specific artifacts.

### Limitations and Considerations

Despite these advances, several limitations warrant acknowledgment: The framework requires paired long-gradient acquisitions for training. While cross-tissue generalization mitigates this requirement, laboratories processing specific sample types may benefit from generating matched training data. Next, our evaluation focused exclusively on the Orbitrap Exploris 480 with Evosep One chromatography. Transferability to alternative mass spectrometers, ion mobility-enabled platforms, or different gradient configurations requires validation, as learned feature patterns may require recalibration when acquisition parameters differ substantially. Instrument-to-instrument variability, even among identical models, can introduce systematic differences in signal characteristics⁴⁴. Sample processing variability in digestion efficiency, peptide recovery, and clean-up may alter spectral features used for classification⁴⁵. Temporal drift in instrument performance, column aging, and calibration state can further introduce heterogeneity between training and inference datasets⁴⁴. Approaches such as batch effect removal neural networks BERNN have been developed to address these challenges in LC-MS classification⁴⁶. Users should evaluate performance on representative samples before large-scale application and consider retraining or batch correction when processing conditions change. Calibration showed conservative behavior at stringent thresholds and modest liberal bias at 10% target RDR in some configurations. Users operating at higher target RDRs should interpret results cautiously and consider orthogonal validation. Finally, our labeling strategy assumes correctness of long-gradient identifications at 1% FDR. Residual false positives in training labels may introduce noise that limits model performance and contributes to calibration imperfections.

### Future Directions

Our implementation did not include variable modifications, but modified peptide candidates carry the same DIA-NN feature set, so the framework should extend to PTM identification. The additional identification challenges posed by modified peptides⁴⁷ suggest rescoring could be particularly useful, though this remains to be validated. Newer instruments such as the Orbitrap Astral routinely identify >10,000 protein groups with shorter gradients⁴⁸, yet the throughput–sensitivity trade-off persists, and testing PeptiDIA on Astral data is a clear next step. Fragment ion predictors (Prosit^20^, MS2PIP⁵⁰) and retention time predictors (DeepLC⁵¹) could provide orthogonal features, and multi-engine rescoring combining DIA-NN with Spectronaut⁵², EncyclopeDIA⁵³, or MSFragger^54^ could improve recovery beyond single-engine approaches. Adapting PeptiDIA to other engines requires changes to feature extraction, but the paired-acquisition training strategy is not engine-specific. Pre-training on large multi-tissue datasets and fine-tuning on limited target data may further reduce the need for paired acquisition training data.

## Conclusion

PeptiDIA is a supervised post-processing framework that leverages DIA-NN output and engineered features to recover peptides under fast-gradient conditions. PeptiDIA requires no modification to acquisition strategies or instrumentation allowing for easy integration into workflows. As DIA proteomics scales to larger cohorts, tools that extract more information from high-throughput acquisitions will become increasingly important. This is particularly relevant in the context of clinical proteomics, where maximizing proteome coverage from short gradients is essential for routine processing of hundreds to thousands of patient samples, as well as for the rapidly emerging field of single-cell proteomics, where ultra-fast acquisition times are required to achieve sufficient cellular throughput. PeptiDIA takes a step in this direction by helping to narrow the gap between speed and sensitivity.

## Acknowledgements

We thank the Proteomics Platform of the Centre de recherche du CHU de Québec - Université Laval for generating the mass spectrometry data used in this study.

## Author Contributions

**Conceptualization**: J.O., M.L., and F.R.-D. **Methodology:** J.O. **Investigation**: J.O. **Formal Analysis**: J.O. **Software**: J.O. **Resources**: B.R. and S.B. **Visualization**: J.O. **Writing – Original Draft**: J.O. **Writing – Review & Editing**: M.L., F.R.-D., and A.D. **Supervision**: M.L., F.R.-D., and A.D. **Funding Acquisition**: A.D. All authors reviewed and approved the final version of the manuscript.

## Data Availability Statement

The mass spectrometry proteomics data for human and murine tissue samples have been deposited to the ProteomeXchange repository under identifier PXD079315. The PeptiDIA source code and pre-trained models are available at https://github.com/Jordano700/PeptiDIA.

## References

(1) Aebersold, R.; Mann, M. Mass Spectrometry-Based Proteomics. Nature 2003, 422 (6928), 198–207. 10.1038/nature01511.

(2) Guo, T.; Steen, J. A.; Mann, M. Mass-Spectrometry-Based Proteomics: From Single Cells to Clinical Applications. Nature 2025, 638 (8052), 901–911. 10.1038/s41586-025-08584-0.

(3) Gillet, L. C.; Navarro, P.; Tate, S.; Röst, H.; Selevsek, N.; Reiter, L.; Bonner, R.; Aebersold, R. Targeted Data Extraction of the MS/MS Spectra Generated by Data-Independent Acquisition: A New Concept for Consistent and Accurate Proteome Analysis. Mol. Cell. Proteomics MCP 2012, 11 (6), O111.016717. 10.1074/mcp.O111.016717.

(4) Lou, R.; Shui, W. Acquisition and Analysis of DIA-Based Proteomic Data: A Comprehensive Survey in 2023. Mol. Cell. Proteomics MCP 2024, 23 (2), 100712. 10.1016/j.mcpro.2024.100712.

(5) Guan, S.; Taylor, P. P.; Han, Z.; Moran, M. F.; Ma, B. Data Dependent–Independent Acquisition (DDIA) Proteomics. J. Proteome Res. 2020, 19 (8), 3230–3237. 10.1021/acs.jproteome.0c00186.

(6) Fröhlich, K.; Fahrner, M.; Brombacher, E.; Seredynska, A.; Maldacker, M.; Kreutz, C.; Schmidt, A.; Schilling, O. Data-Independent Acquisition: A Milestone and Prospect in Clinical Mass Spectrometry-Based Proteomics. Mol. Cell. Proteomics MCP 2024, 23 (8), 100800. 10.1016/j.mcpro.2024.100800.

(7) Lenčo, J.; Jadeja, S.; Naplekov, D. K.; Krokhin, O. V.; Khalikova, M. A.; Chocholouš, P.; Urban, J.; Broeckhoven, K.; Nováková, L.; Švec, F. Reversed-Phase Liquid Chromatography of Peptides for Bottom-Up Proteomics: A Tutorial. J. Proteome Res. 2022, 21 (12), 2846–2892. 10.1021/acs.jproteome.2c00407.

(8) Neue, U. D. Theory of Peak Capacity in Gradient Elution. J. Chromatogr. A 2005, 1079(1–2), 153–161. 10.1016/j.chroma.2005.03.008.

(9) Hsieh, E. J.; Bereman, M. S.; Durand, S.; Valaskovic, G. A.; MacCoss, M. J. Effects of Column and Gradient Lengths on Peak Capacity and Peptide Identification in Nanoflow LC-MS/MS of Complex Proteomic Samples. J. Am. Soc. Mass Spectrom. 2013, 24 (1), 148–153. 10.1007/s13361-012-0508-6.

(10) Bekker-Jensen, D. B.; Martínez-Val, A.; Steigerwald, S.; Rüther, P.; Fort, K. L.; Arrey, T. N.; Harder, A.; Makarov, A.; Olsen, J. V. A Compact Quadrupole-Orbitrap Mass Spectrometer with FAIMS Interface Improves Proteome Coverage in Short LC Gradients. Mol. Cell. Proteomics MCP 2020, 19 (4), 716–729. 10.1074/mcp.TIR119.001906.

(11) Kawashima, Y.; Nagai, H.; Konno, R.; Ishikawa, M.; Nakajima, D.; Sato, H.; Nakamura, R.; Furuyashiki, T.; Ohara, O. Single-Shot 10K Proteome Approach: Over 10,000 Protein Identifications by Data-Independent Acquisition-Based Single-Shot Proteomics with Ion Mobility Spectrometry. J. Proteome Res. 2022, 21 (6), 1418–1427. 10.1021/acs.jproteome.2c00023.

(12) Messner, C. B.; Demichev, V.; Wendisch, D.; Michalick, L.; White, M.; Freiwald, A.; Textoris-Taube, K.; Vernardis, S. I.; Egger, A.-S.; Kreidl, M.; Ludwig, D.; Kilian, C.; Agostini, F.; Zelezniak, A.; Thibeault, C.; Pfeiffer, M.; Hippenstiel, S.; Hocke, A.; von Kalle, C.; Campbell, A.; Hayward, C.; Porteous, D. J.; Marioni, R. E.; Langenberg, C.; Lilley, K. S.; Kuebler, W. M.; Mülleder, M.; Drosten, C.; Suttorp, N.; Witzenrath, M.; Kurth, F.; Sander, L. E.; Ralser, M. Ultra-High-Throughput Clinical Proteomics Reveals Classifiers of COVID-19 Infection. Cell Syst. 2020, 11 (1), 11–24.e4. 10.1016/j.cels.2020.05.012.

(13) Messner, C. B.; Demichev, V.; Bloomfield, N.; Yu, J. S. L.; White, M.; Kreidl, M.; Egger, A.-S.; Freiwald, A.; Ivosev, G.; Wasim, F.; Zelezniak, A.; Jürgens, L.; Suttorp, N.; Sander, L. E.; Kurth, F.; Lilley, K. S.; Mülleder, M.; Tate, S.; Ralser, M. Ultra-Fast Proteomics with Scanning SWATH. Nat. Biotechnol. 2021, 39 (7), 846–854. 10.1038/s41587-021-00860-4.

(14) Bache, N.; Geyer, P. E.; Bekker-Jensen, D. B.; Hoerning, O.; Falkenby, L.; Treit, P. V.; Doll, S.; Paron, I.; Müller, J. B.; Meier, F.; Olsen, J. V.; Vorm, O.; Mann, M. A Novel LC System Embeds Analytes in Pre-Formed Gradients for Rapid, Ultra-Robust Proteomics. Mol. Cell. Proteomics MCP 2018, 17 (11), 2284–2296. 10.1074/mcp.TIR118.000853.

(15) Meier, F.; Brunner, A.-D.; Frank, M.; Ha, A.; Bludau, I.; Voytik, E.; Kaspar-Schoenefeld, S.; Lubeck, M.; Raether, O.; Bache, N.; Aebersold, R.; Collins, B. C.; Röst, H. L.; Mann, M. diaPASEF: Parallel Accumulation–Serial Fragmentation Combined with Data-Independent Acquisition. Nat. Methods 2020, 17 (12), 1229–1236. 10.1038/s41592-020-00998-0.

(16) Beck, A. G.; Muhoberac, M.; Randolph, C. E.; Beveridge, C. H.; Wijewardhane, P. R.; Kenttämaa, H. I.; Chopra, G. Recent Developments in Machine Learning for Mass Spectrometry. ACS Meas. Sci. Au 2024, 4 (3), 233–246. 10.1021/acsmeasuresciau.3c00060.

(17) Che, Y.; Zhao, M.; Gao, Y.; Zhang, Z.; Zhang, X. Application of Machine Learning for Mass Spectrometry-Based Multi-Omics in Thyroid Diseases. Front. Mol. Biosci. 2024, 11, 1483326. 10.3389/fmolb.2024.1483326.

(18) Coyle, E.; Leclercq, M.; Gotti, C.; Roux-Dalvai, F.; Droit, A. ProPickML: Advancing Clinical Diagnostics with Automated Peak Picking in Label-Free Targeted Proteomics. J. Proteome Res. 2025, 24 (1), 244–255. 10.1021/acs.jproteome.4c00689.

(19) Käll, L.; Canterbury, J. D.; Weston, J.; Noble, W. S.; MacCoss, M. J. Semi-SupervisedLearning for Peptide Identification from Shotgun Proteomics Datasets. Nat. Methods 2007, 4 (11), 923–925. 10.1038/nmeth1113.

(20) Gessulat, S.; Schmidt, T.; Zolg, D. P.; Samaras, P.; Schnatbaum, K.; Zerweck, J.; Knaute, T.; Rechenberger, J.; Delanghe, B.; Huhmer, A.; Reimer, U.; Ehrlich, H.-C.; Aiche, S.; Kuster, B.; Wilhelm, M. Prosit: Proteome-Wide Prediction of Peptide Tandem Mass Spectra by Deep Learning. Nat. Methods 2019, 16 (6), 509–518. 10.1038/s41592-019-0426-7.

(21) Meier, F.; Köhler, N. D.; Brunner, A.-D.; Wanka, J.-M. H.; Voytik, E.; Strauss, M. T.; Theis, F. J.; Mann, M. Deep Learning the Collisional Cross Sections of the Peptide Universe from a Million Experimental Values. Nat. Commun. 2021, 12 (1), 1185. 10.1038/s41467-021-21352-8.

(22) Mann, M.; Kumar, C.; Zeng, W.-F.; Strauss, M. T. Artificial Intelligence for Proteomics and Biomarker Discovery. Cell Syst. 2021, 12 (8), 759–770. 10.1016/j.cels.2021.06.006.

(23) Demichev, V.; Messner, C. B.; Vernardis, S. I.; Lilley, K. S.; Ralser, M. DIA-NN: Neural Networks and Interference Correction Enable Deep Proteome Coverage in High Throughput. Nat. Methods 2020, 17 (1), 41–44. 10.1038/s41592-019-0638-x.

(24) Doellinger, J.; Blumenscheit, C.; Schneider, A.; Lasch, P. Increasing Proteome Depth While Maintaining Quantitative Precision in Short-Gradient Data-Independent Acquisition Proteomics. J. Proteome Res. 2023, 22 (6), 2131–2140. 10.1021/acs.jproteome.3c00078.

(25) Martens, L.; Chambers, M.; Sturm, M.; Kessner, D.; Levander, F.; Shofstahl, J.; Tang, W. H.; Römpp, A.; Neumann, S.; Pizarro, A. D.; Montecchi-Palazzi, L.; Tasman, N.; Coleman, M.; Reisinger, F.; Souda, P.; Hermjakob, H.; Binz, P.-A.; Deutsch, E. W. mzML—a Community Standard for Mass Spectrometry Data*. Mol. Cell. Proteomics 2011, 10 (1), R110.000133. 10.1074/mcp.R110.000133.

(26) Hulstaert, N.; Shofstahl, J.; Sachsenberg, T.; Walzer, M.; Barsnes, H.; Martens, L.; Perez-Riverol, Y. ThermoRawFileParser: Modular, Scalable, and Cross-Platform RAW File Conversion. J. Proteome Res. 2020, 19 (1), 537–542. 10.1021/acs.jproteome.9b00328.

(27) UniProt. UniProt. https://www.uniprot.org/proteomes/UP000005640 (accessed 2025-11-09).

(28) UniProt. UniProt. https://www.uniprot.org/proteomes/UP000000589 (accessed 2025-11-09).

(29) Chen, T.; Guestrin, C. XGBoost: A Scalable Tree Boosting System. In Proceedings of the 22nd ACM SIGKDD International Conference on Knowledge Discovery and Data Mining; KDD ‘16; Association for Computing Machinery: New York, NY, USA, 2016; pp 785–794. 10.1145/2939672.2939785.

(30) Akiba, T.; Sano, S.; Yanase, T.; Ohta, T.; Koyama, M. Optuna: A Next-Generation Hyperparameter Optimization Framework. In Proceedings of the 25th ACM SIGKDD International Conference on Knowledge Discovery & Data Mining; KDD ‘19; Association for Computing Machinery: New York, NY, USA, 2019; pp 2623–2631. 10.1145/3292500.3330701.

(31) Fawcett, T. An Introduction to ROC Analysis. Pattern Recognit. Lett. 2006, 27 (8), 861–874. 10.1016/j.patrec.2005.10.010.

(32) Zadrozny, B.; Elkan, C. Transforming Classifier Scores into Accurate Multiclass Probability Estimates. In Proceedings of the eighth ACM SIGKDD international conference on Knowledge discovery and data mining; KDD ‘02; Association for Computing Machinery: New York, NY, USA, 2002; pp 694–699. 10.1145/775047.775151.

(33) Niculescu-Mizil, A.; Caruana, R. Predicting Good Probabilities with Supervised Learning. In Proceedings of the 22nd international conference on Machine learning; ICML ‘05; Association for Computing Machinery: New York, NY, USA, 2005; pp 625–632. 10.1145/1102351.1102430.

(34) Spearman, C. The Proof and Measurement of Association between Two Things. Am. J. Psychol. 1904, 15 (1), 72–101. 10.2307/1412159.

(35) Mann, H. B.; Whitney, D. R. On a Test of Whether One of Two Random Variables Is Stochastically Larger than the Other. Ann. Math. Stat. 1947, 18 (1), 50–60. 10.1214/aoms/1177730491.

(36) Lundberg, S. M.; Lee, S.-I. A Unified Approach to Interpreting Model Predictions. In Proceedings of the 31st International Conference on Neural Information Processing Systems; NIPS’17; Curran Associates Inc.: Red Hook, NY, USA, 2017; pp 4768–4777.

(37) Scikit-learn: Machine Learning in Python | The Journal of Machine Learning Research. https://dl.acm.org/doi/10.5555/1953048.2078195 (accessed 2026-01-12).

(38) McKinney, W. Data Structures for Statistical Computing in Python; Austin, Texas, 2010; pp 56–61. 10.25080/Majora-92bf1922-00a.

(39) Harris, C. R.; Millman, K. J.; van der Walt, S. J.; Gommers, R.; Virtanen, P.; Cournapeau, D.; Wieser, E.; Taylor, J.; Berg, S.; Smith, N. J.; Kern, R.; Picus, M.; Hoyer, S.; van Kerkwijk, M. H.; Brett, M.; Haldane, A.; Del Río, J. F.; Wiebe, M.; Peterson, P.; Gérard-Marchant, P.; Sheppard, K.; Reddy, T.; Weckesser, W.; Abbasi, H.; Gohlke, C.; Oliphant, T. E. Array Programming with NumPy. Nature 2020, 585 (7825), 357–362. 10.1038/s41586-020-2649-2.

(40) Streamlit • A faster way to build and share data apps. https://streamlit.io/ (accessed 2026-01-12).

(41) Du, P.; Stolovitzky, G.; Horvatovich, P.; Bischoff, R.; Lim, J.; Suits, F. A Noise Model for Mass Spectrometry Based Proteomics. Bioinformatics 2008, 24 (8), 1070–1077. 10.1093/bioinformatics/btn078.

(42) Liigand, P.; Kaupmees, K.; Kruve, A. Influence of the Amino Acid Composition on the Ionization Efficiencies of Small Peptides. J. Mass Spectrom. JMS 2019, 54 (6), 481–487. 10.1002/jms.4348.

(43) Bantscheff, M.; Schirle, M.; Sweetman, G.; Rick, J.; Kuster, B. Quantitative Mass Spectrometry in Proteomics: A Critical Review. Anal. Bioanal. Chem. 2007, 389 (4), 1017–1031. 10.1007/s00216-007-1486-6.

(44) Èuklina, J.; Lee, C. H.; Williams, E. G.; Sajic, T.; Collins, B. C.; Rodríguez Martínez, M.; Sharma, V. S.; Wendt, F.; Goetze, S.; Keele, G. R.; Wollscheid, B.; Aebersold, R.; Pedrioli, P. G. A. Diagnostics and Correction of Batch Effects in Large-Scale Proteomic Studies: A Tutorial. Mol. Syst. Biol. 2021, 17 (8), e10240. 10.15252/msb.202110240.

(45) Piehowski, P. D.; Petyuk, V. A.; Orton, D. J.; Xie, F.; Moore, R. J.; Ramirez-Restrepo, M.; Engel, A.; Lieberman, A. P.; Albin, R. L.; Camp, D. G.; Smith, R. D.; Myers, A. J. Sources of Technical Variability in Quantitative LC-MS Proteomics: Human Brain Tissue Sample Analysis. J. Proteome Res. 2013, 12 (5), 2128–2137. 10.1021/pr301146m.

(46) Pelletier, S. J.; Leclercq, M.; Roux-Dalvai, F.; de Geus, M. B.; Leslie, S.; Wang, W.; Lam, T. T.; Nairn, A. C.; Arnold, S. E.; Carlyle, B. C.; Precioso, F.; Droit, A. BERNN: Enhancing Classification of Liquid Chromatography Mass Spectrometry Data with Batch Effect Removal Neural Networks. Nat. Commun. 2024, 15 (1), 3777. 10.1038/s41467-024-48177-5.

(47) Solari, F. A.; Dell’Aica, M.; Sickmann, A.; Zahedi, R. P. Why Phosphoproteomics Is Still a Challenge. Mol. Biosyst. 2015, 11 (6), 1487–1493. 10.1039/c5mb00024f.

(48) Guzman, U. H.; Martinez-Val, A.; Ye, Z.; Damoc, E.; Arrey, T. N.; Pashkova, A.; Renuse, S.; Denisov, E.; Petzoldt, J.; Peterson, A. C.; Harking, F.; Østergaard, O.; Rydbirk, R.; Aznar, S.; Stewart, H.; Xuan, Y.; Hermanson, D.; Horning, S.; Hock, C.; Makarov, A.; Zabrouskov, V.; Olsen, J. V. Ultra-Fast Label-Free Quantification and Comprehensive Proteome Coverage with Narrow-Window Data-Independent Acquisition. Nat. Biotechnol. 2024, 42 (12), 1855–1866. 10.1038/s41587-023-02099-7.

(49) Declercq, A.; Bouwmeester, R.; Chiva, C.; Sabidó, E.; Hirschler, A.; Carapito, C.; Martens, L.; Degroeve, S.; Gabriels, R. Updated MS^2^PIP Web Server Supports Cutting-Edge Proteomics Applications. Nucleic Acids Res. 2023, 51(), W338–W342. 10.1093/nar/gkad335.

(50) Bouwmeester, R.; Gabriels, R.; Hulstaert, N.; Martens, L.; Degroeve, S. DeepLC Can Predict Retention Times for Peptides That Carry As-yet Unseen Modifications. Nat. Methods 2021, 18 (11), 1363–1369. 10.1038/s41592-021-01301-5.

(51) Bruderer, R.; Bernhardt, O. M.; Gandhi, T.; Miladinović, S. M.; Cheng, L.-Y.; Messner, S.; Ehrenberger, T.; Zanotelli, V.; Butscheid, Y.; Escher, C.; Vitek, O.; Rinner, O.; Reiter, L. Extending the Limits of Quantitative Proteome Profiling with Data-Independent Acquisition and Application to Acetaminophen-Treated Three-Dimensional Liver Microtissues. Mol. Cell. Proteomics MCP 2015, 14 (5), 1400–1410. 10.1074/mcp.M114.044305.

(52) Searle, B. C.; Pino, L. K.; Egertson, J. D.; Ting, Y. S.; Lawrence, R. T.; MacLean, B. X.; Villén, J.; MacCoss, M. J. Chromatogram Libraries Improve Peptide Detection and Quantification by Data Independent Acquisition Mass Spectrometry. Nat. Commun. 2018, 9 (1), 5128. 10.1038/s41467-018-07454-w.

(53) Kong, A. T.; Leprevost, F. V.; Avtonomov, D. M.; Mellacheruvu, D.; Nesvizhskii, A. I. MSFragger: Ultrafast and Comprehensive Peptide Identification in Mass Spectrometry–Based Proteomics. Nat. Methods 2017, 14 (5), 513–520. 10.1038/nmeth.4256.

