## Supplementary Methods for "PeptiDIA: A Machine Learning Framework for Enhanced Peptide Identification in Fast-Gradient Data-Independent Acquisition Proteomics"

### Sample Preparation

Three protocols were used depending on tissue type, as described below.

#### Protocol 1 (Ileum, Colon and Liver Tissues)

- Frozen tissues were subjected to bead beating and sonication in 50 mM ammonium bicarbonate (ABC), 0.5% sodium deoxycholate (SDC), 50 mM dithiothreitol (DTT), protease inhibitors (Roche Complete Mini, pepstatin 1 µM).
- After centrifugation to remove debris, proteins were precipitated by addition of ice-cold acetone overnight.
- After centrifugation, the supernatant was discarded and the pellet resuspended in 50 mM ABC / 0.5% DOC.
- 20 µg of protein (based on Bradford assay) were heated at 95°C for 5 min for protein denaturation
- Cysteine disulfide bonds reduction and alkylation were carried out by addition of 0.2 mM DTT (final concentration) for 30 min, 37°C followed by addition of 0.8 mM iodoacetamide (final concentration) for 30 min, 37° in the dark.
- Digestion was performed with 400 ng trypsin (Promega), overnight at 37°C and then stopped by addition of formic acid (FA).
- Samples were centrifuged and supernatant vacuum-dried prior to resuspension in 0.1% FA.

#### Protocol 2 (Coronary Artery and Epicardial Adipose Tissues)

- Proteins in Laemmli sample buffer were loaded on an SDS-PAGE gel. The protein migration was stopped at the interface between stacking and running parts of the gel to collect the whole protein extract. After Coomassie staining, a unique gel band containing all the proteins of each sample was excised.
- Cysteine disulfide bonds were reduces in gel with 10 mM DTT and alkylate in gel with 55 mM iodoacetamide.
- In gel digestion was performed with 126 nM modified porcine trypsin (Promega) at 37°C. overnight
- Resulting peptides were sequentially extracted with: 1% formic acid / 2% acetonitrile (ACN), then 1% formic acid / 50% CAN, then pooled and vacuum dried.
- The samples were resuspended in 0.1% FA and peptide concentration was determined by absorbance measurement at 205nm.

#### Protocol 3 (Heart Tissue)

- Tissues were cryogenically pulverized using a CryoMill and resuspended in 5% SDS / 50 mM TEAB, pH 8.5, with Complete Mini EDTA-free protease inhibitor (Roche) prior to sonication with a Bioruptor.
- Sample were centrifugated and the protein concentration in supernatant was determined using Pierce BCA Protein Assay.
- Protein denaturation, cysteine disulfide bonds reduction and alkylation and protein digestion on S-Trap (ProtiFi) was performed according to manufacturer protocol.
- The vacuum dried resulting peptides were resuspended in 0.1% FA and peptide concentration was determined using Pierce Quantitative Fluorometric Peptide Assay.

### LC-MS/MS analysis

All tissues were analyzed using the same two methods (30SPD and 300SPD)

- For each LC-MS/MS analysis, 250ng of peptides were loading on Evotips Pure (Evosep) according to manufacturer protocol and injected on an Evosep One liquid chromatography system (Evosep) interfaced with an Orbitrap Exploris 480 mass spectrometer (Thermo Fisher Scientific).

- Two manufacturer pre-programmed gradients were used: 30 or 300 Samples Per Day (SPD). A 15 cm length x 150 µm ID C18 column (EV1137) and a 4 cm length x 150 µm ID C18 column (EV1107) were used for 30 and 300SPD gradients respectively.

- The mass spectrometer was operating in DIA mode with the following DIA schemes and parameters:

For 30 SPD gradients: 1x MS1 scan (Mass range: 350-1500m/z; Orbitrap resolution: 60,000; AGC target: 300%, Max Injection Time: 118ms) and 42 x MS2 scans (Isolation window: 13m/z, Precursor mass range: 350-896; HCD: 30%; Orbitrap resolution: 22,500; AGC target: 800%, Max Injection Time: 38ms)
For 300 SPD gradients: 1x MS1 scan (Mass range: 350-1500m/z; Orbitrap resolution: 30,000; AGC target: 300%, Max Injection Time: 54ms) and 21 x MS2 scans (Isolation window: 25m/z, Precursor mass range: 350-875; HCD: 30%; Orbitrap resolution: 7,500; AGC target: 800%, Max Injection Time: 6ms)
