## Supplementary Data for "PeptiDIA: A Machine Learning Framework for Enhanced Peptide Identification in Fast-Gradient Data-Independent Acquisition Proteomics"

**Table S1. Input features for the PeptiDIA machine learning model**. A total of 87 features is provided to the XGBoost classifier, which performs internal feature selection during training based on information gain.

| No. | Feature | Description | Source | Category |
|---|---|---|---|---|
| 1 | **GG.Q.Value** | Gene group q-value (primary FDR metric) | DIA-NN | Quality Metrics |
| 2 | **Q.Value** | Precursor-level q-value | DIA-NN | Quality Metrics |
| 3 | **PEP** | Posterior Error Probability | DIA-NN | Quality Metrics |
| 4 | **PG.Q.Value** | Protein group q-value | DIA-NN | Quality Metrics |
| 5 | **PG.PEP** | Protein group posterior error probability | DIA-NN | Quality Metrics |
| 6 | **Global.Q.Value** | Global q-value across all runs | DIA-NN | Quality Metrics |
| 7 | **Global.PG.Q.Value** | Global protein group q-value | DIA-NN | Quality Metrics |
| 8 | **Global.Peptidoform.Q.Value** | Global peptidoform q-value | DIA-NN | Quality Metrics |
| 9 | **Protein.Q.Value** | Protein-level q-value | DIA-NN | Quality Metrics |
| 10 | **Quantity.Quality** | Quantification reliability score (0-1) | DIA-NN | Quality Metrics |
| 11 | **PG.MaxLFQ.Quality** | Protein group MaxLFQ quality | DIA-NN | Quality Metrics |
| 12 | **Genes.MaxLFQ.Quality** | Gene-level MaxLFQ quality | DIA-NN | Quality Metrics |
| 13 | **Genes.MaxLFQ.Unique.Quality** | Unique gene-level MaxLFQ quality | DIA-NN | Quality Metrics |
| 14 | **Proteotypic** | Proteotypic peptide indicator (0/1) | DIA-NN | Quality Metrics |
| 15 | **Evidence** | Number of supporting evidence points | DIA-NN | Quality Metrics |
| 16 | **Mass.Evidence** | Mass-based evidence count | DIA-NN | Quality Metrics |
| 17 | **Channel.Evidence** | Channel-based evidence count | DIA-NN | Quality Metrics |
| 18 | **PG.MaxLFQ** | Protein group MaxLFQ intensity | DIA-NN | Quality Metrics |
| 19 | **Genes.MaxLFQ** | Gene-level MaxLFQ intensity | DIA-NN | Quality Metrics |
| 20 | **Genes.MaxLFQ.Unique** | Unique peptide gene-level MaxLFQ | DIA-NN | Quality Metrics |
| 21 | **RT** | Observed retention time (minutes) | DIA-NN | MS/Chromatography |
| 22 | **iRT** | Indexed retention time (normalized) | DIA-NN | MS/Chromatography |
| 23 | **Predicted.RT** | Deep learning predicted RT | DIA-NN | MS/Chromatography |
| 24 | **Predicted.iRT** | Predicted indexed RT | DIA-NN | MS/Chromatography |
| 25 | **RT.Start** | RT window start | DIA-NN | MS/Chromatography |
| 26 | **RT.Stop** | RT window end | DIA-NN | MS/Chromatography |
| 27 | **Precursor.Mz** | Precursor mass-to-charge ratio | DIA-NN | MS/Chromatography |
| 28 | **Precursor.Charge** | Precursor charge state | DIA-NN | MS/Chromatography |
| 29 | **Best.Fr.Mz** | Best fragment m/z | DIA-NN | MS/Chromatography |
| 30 | **Best.Fr.Mz.Delta** | Best fragment mass error (ppm) | DIA-NN | MS/Chromatography |
| 31 | **Ms1.Apex.Mz.Delta** | MS1 apex mass error (ppm) | DIA-NN | MS/Chromatography |
| 32 | **Ms1.Area** | MS1 extracted ion chromatogram area | DIA-NN | MS/Chromatography |
| 33 | **Ms1.Apex.Area** | MS1 apex peak area | DIA-NN | MS/Chromatography |
| 34 | **Precursor.Quantity** | Precursor quantification | DIA-NN | MS/Chromatography |
| 35 | **Precursor.Normalised** | Normalized precursor intensity | DIA-NN | MS/Chromatography |
| 36 | **Ms1.Normalised** | Normalized MS1 intensity | DIA-NN | MS/Chromatography |
| 37 | **FWHM** | Full width at half maximum | DIA-NN | MS/Chromatography |

| No. | Feature | Description | Source | Category |
|---|---|---|---|---|
| 38 | **Ms1.Profile.Corr** | MS1 isotope profile correlation | DIA-NN | MS/Chromatography |
| 39 | **Ms1.Total.Signal.Before** | Total MS1 signal before apex | DIA-NN | MS/Chromatography |
| 40 | **Ms1.Total.Signal.After** | Total MS1 signal after apex | DIA-NN | MS/Chromatography |
| 41 | **iIM** | Indexed ion mobility | DIA-NN | MS/Chromatography |
| 42 | **Precursor.Lib.Index** | Library precursor index | DIA-NN | MS/Chromatography |
| 43 | **sequence_length** | Total amino acid count in peptide | Engineered | Sequence |
| 44 | **aa_count_A** | Count of Alanine (A) residues | Engineered | Sequence |
| 45 | **aa_count_C** | Count of Cysteine (C) residues | Engineered | Sequence |
| 46 | **aa_count_D** | Count of Aspartic acid (D) residues | Engineered | Sequence |
| 47 | **aa_count_E** | Count of Glutamic acid (E) residues | Engineered | Sequence |
| 48 | **aa_count_F** | Count of Phenylalanine (F) residues | Engineered | Sequence |
| 49 | **aa_count_G** | Count of Glycine (G) residues | Engineered | Sequence |
| 50 | **aa_count_H** | Count of Histidine (H) residues | Engineered | Sequence |
| 51 | **aa_count_I** | Count of Isoleucine (I) residues | Engineered | Sequence |
| 52 | **aa_count_K** | Count of Lysine (K) residues | Engineered | Sequence |
| 53 | **aa_count_L** | Count of Leucine (L) residues | Engineered | Sequence |
| 54 | **aa_count_M** | Count of Methionine (M) residues | Engineered | Sequence |
| 55 | **aa_count_N** | Count of Asparagine (N) residues | Engineered | Sequence |
| 56 | **aa_count_P** | Count of Proline (P) residues | Engineered | Sequence |
| 57 | **aa_count_Q** | Count of Glutamine (Q) residues | Engineered | Sequence |
| 58 | **aa_count_R** | Count of Arginine (R) residues | Engineered | Sequence |
| 59 | **aa_count_S** | Count of Serine (S) residues | Engineered | Sequence |
| 60 | **aa_count_T** | Count of Threonine (T) residues | Engineered | Sequence |
| 61 | **aa_count_V** | Count of Valine (V) residues | Engineered | Sequence |
| 62 | **aa_count_W** | Count of Tryptophan (W) residues | Engineered | Sequence |
| 63 | **aa_count_Y** | Count of Tyrosine (Y) residues | Engineered | Sequence |
| 64 | **aa_freq_A** | Frequency of Alanine (A) | Engineered | Sequence |
| 65 | **aa_freq_C** | Frequency of Cysteine (C) | Engineered | Sequence |
| 66 | **aa_freq_D** | Frequency of Aspartic acid (D) | Engineered | Sequence |
| 67 | **aa_freq_E** | Frequency of Glutamic acid (E) | Engineered | Sequence |
| 68 | **aa_freq_F** | Frequency of Phenylalanine (F) | Engineered | Sequence |
| 69 | **aa_freq_G** | Frequency of Glycine (G) | Engineered | Sequence |
| 70 | **aa_freq_H** | Frequency of Histidine (H) | Engineered | Sequence |
| 71 | **aa_freq_I** | Frequency of Isoleucine (I) | Engineered | Sequence |
| 72 | **aa_freq_K** | Frequency of Lysine (K) | Engineered | Sequence |
| 73 | **aa_freq_L** | Frequency of Leucine (L) | Engineered | Sequence |
| 74 | **aa_freq_M** | Frequency of Methionine (M) | Engineered | Sequence |
| 75 | **aa_freq_N** | Frequency of Asparagine (N) | Engineered | Sequence |
| 76 | **aa_freq_P** | Frequency of Proline (P) | Engineered | Sequence |
| 77 | **aa_freq_Q** | Frequency of Glutamine (Q) | Engineered | Sequence |
| 78 | **aa_freq_R** | Frequency of Arginine (R) | Engineered | Sequence |
| 79 | **aa_freq_S** | Frequency of Serine (S) | Engineered | Sequence |

| No. | Feature | Description | Source | Category |
|---|---|---|---|---|
| 80 | **aa_freq_T** | Frequency of Threonine (T) | Engineered | Sequence |
| 81 | **aa_freq_V** | Frequency of Valine (V) | Engineered | Sequence |
| 82 | **aa_freq_W** | Frequency of Tryptophan (W) | Engineered | Sequence |
| 83 | **aa_freq_Y** | Frequency of Tyrosine (Y) | Engineered | Sequence |
| 84 | **log_Ms1.Area** | $\log_2$(Ms1.Area) | Engineered | Derived |
| 85 | **log_Precursor.Quantity** | $\log_2$(Precursor.Quantity) | Engineered | Derived |
| 86 | **zscore_Ms1.Area** | Z-score normalized Ms1.Area | Engineered | Derived |
| 87 | **zscore_Precursor.Quantity** | Z-score normalized Precursor.Quantity | Engineered | Derived |

**Table S2. XGBoost hyperparameters.** Values were determined by grid search optimization across all tissue datasets.

| No. | Hyperparameter | Value | Description |
|---|---|---|---|
| 1 | **n_estimators** | 600 | Number of boosting rounds |
| 2 | **max_depth** | 8 | Maximum tree depth |
| 3 | **learning_rate** | 0.1 | Step size shrinkage per boosting round |
| 4 | **subsample** | 0.8 | Fraction of samples used per tree |
| 5 | **colsample_bytree** | 0.6 | Fraction of features used per tree |
| 6 | **reg_lambda** | 1.5 | L2 regularization term on weights |
| 7 | **min_child_weight** | 3 | Minimum sum of instance weight in a child node |

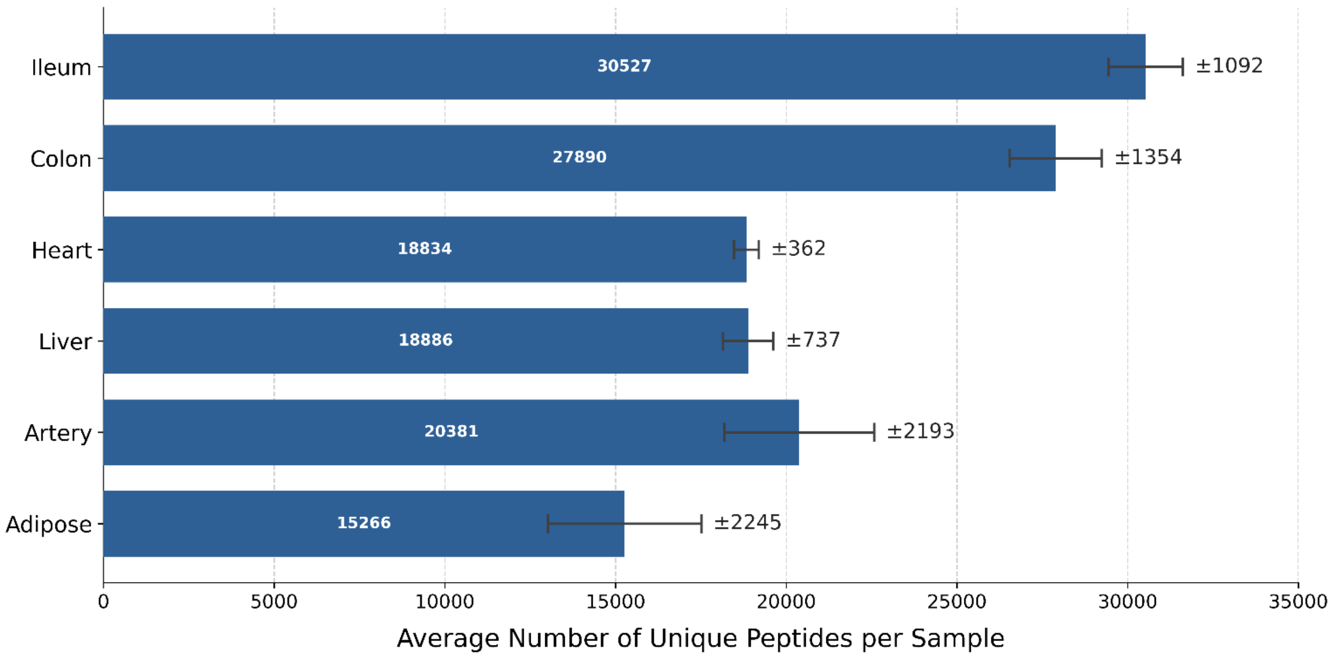


**Figure S1. Average number of unique peptides per sample in long-gradient 30SPD runs (DIA-NN at 1% FDR) across six tissue types.** Bars show mean unique modified sequences per tissue and error bars indicate standard deviation across samples.

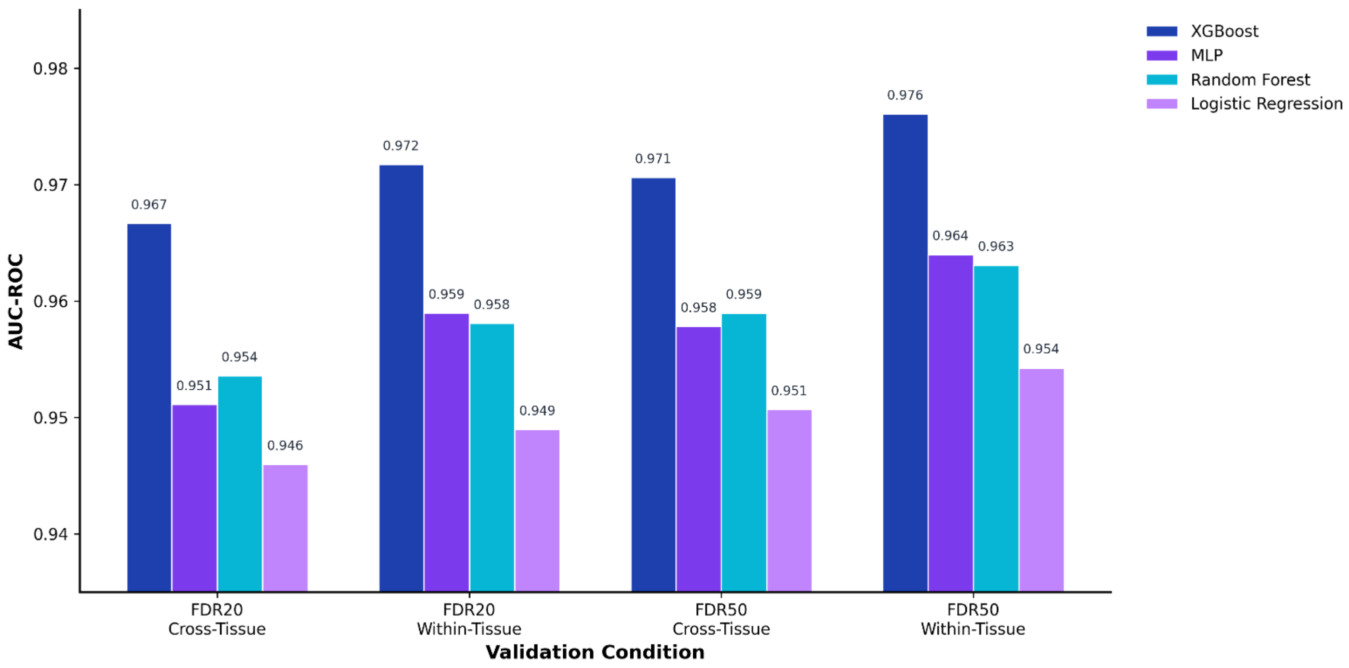


**Figure S2. Comparison of machine learning models for peptide validation across validation conditions.** Four models were evaluated: XGBoost, Random Forest, Multi-Layer Perceptron (MLP), and Logistic Regression. Performance was assessed using Area Under the Receiver Operating Characteristic curve (AUC-ROC), a threshold-independent metric that enables fair comparison without requiring optimal probability calibration for each model. All model hyperparameters were optimized using Optuna with 30 trials per model to ensure fair comparison. Models were assessed under four conditions combining two FDR training pools (FDR20/50) and two validation strategies (cross-tissue: leave-one-tissue-out; within-tissue: leave-one-sample-out).

### Table S3. Peptide and Protein Group Percentage Increases Across Training Strategies

#### (A) FDR20 Cross-Tissue

| Tissue | Target RDR | Peptide Increase (%) | Protein Increase (%) |
|---|---|---|---|
| Artery | 1% | 40.7 | 24.0 |
| Artery | 2% | 47.1 | 24.1 |
| Artery | 5% | 52.8 | 24.4 |
| Artery | 10% | 55.5 | 24.4 |
| Heart | 1% | 21.5 | 9.9 |
| Heart | 2% | 26.0 | 10.3 |
| Heart | 5% | 30.5 | 10.5 |
| Heart | 10% | 32.7 | 10.7 |
| Colon | 1% | 23.8 | 16.2 |
| Colon | 2% | 29.7 | 16.9 |
| Colon | 5% | 35.0 | 17.1 |
| Colon | 10% | 37.3 | 17.2 |
| Liver | 1% | 17.5 | 11.6 |
| Liver | 2% | 21.7 | 12.5 |
| Liver | 5% | 25.6 | 12.9 |
| Liver | 10% | 27.4 | 13.0 |
| Adipose | 1% | 37.1 | 19.4 |
| Adipose | 2% | 45.3 | 20.8 |
| Adipose | 5% | 49.5 | 21.4 |
| Adipose | 10% | 52.0 | 21.6 |
| Ileum | 1% | 26.6 | 17.1 |
| Ileum | 2% | 31.3 | 17.4 |
| Ileum | 5% | 36.9 | 17.7 |
| Ileum | 10% | 39.4 | 17.8 |
| **Mean** | 1% | 27.9 | 16.4 |
| **Mean** | 2% | 33.5 | 17.0 |
| **Mean** | 5% | 38.4 | 17.3 |
| **Mean** | 10% | 40.7 | 17.4 |

#### (B) FDR20 Within-Tissue

| Tissue | Target RDR | Peptide Increase (%) | Protein Increase (%) |
|---|---|---|---|
| Artery | 1% | 36.5 | 23.5 |
| Artery | 2% | 46.1 | 24.1 |
| Artery | 5% | 53.5 | 24.4 |
| Artery | 10% | 56.4 | 24.5 |
| Heart | 1% | 30.3 | 10.5 |
| Heart | 2% | 32.1 | 10.6 |
| Heart | 5% | 34.0 | 10.7 |
| Heart | 10% | 34.9 | 10.8 |
| Colon | 1% | 24.8 | 16.6 |
| Colon | 2% | 29.4 | 16.9 |
| Colon | 5% | 35.0 | 17.1 |
| Colon | 10% | 37.5 | 17.2 |
| Liver | 1% | 21.0 | 12.5 |
| Liver | 2% | 23.6 | 12.8 |
| Liver | 5% | 26.9 | 13.0 |
| Liver | 10% | 28.2 | 13.0 |
| Adipose | 1% | 39.3 | 20.2 |
| Adipose | 2% | 47.7 | 21.2 |
| Adipose | 5% | 51.7 | 21.5 |
| Adipose | 10% | 53.6 | 21.6 |
| Ileum | 1% | 28.1 | 17.3 |
| Ileum | 2% | 32.9 | 17.5 |
| Ileum | 5% | 37.8 | 17.7 |
| Ileum | 10% | 40.1 | 17.8 |
| **Mean** | 1% | 30.0 | 16.8 |
| **Mean** | 2% | 35.3 | 17.2 |
| **Mean** | 5% | 39.8 | 17.4 |
| **Mean** | 10% | 41.8 | 17.5 |

#### (C) FDR50 Cross-Tissue

| Tissue | Target RDR | Peptide Increase (%) | Protein Increase (%) |
|---|---|---|---|
| Artery | 1% | 42.2 | 23.7 |
| Artery | 2% | 53.1 | 24.3 |
| Artery | 5% | 61.4 | 24.5 |
| Artery | 10% | 65.6 | 24.7 |
| Heart | 1% | 16.0 | 8.0 |
| Heart | 2% | 27.7 | 10.3 |
| Heart | 5% | 36.0 | 11.0 |
| Heart | 10% | 39.3 | 11.2 |
| Colon | 1% | 21.3 | 15.4 |
| Colon | 2% | 31.9 | 17.1 |
| Colon | 5% | 39.8 | 17.4 |
| Colon | 10% | 45.1 | 17.5 |
| Liver | 1% | 16.5 | 10.6 |
| Liver | 2% | 23.2 | 12.7 |
| Liver | 5% | 29.7 | 13.0 |
| Liver | 10% | 32.6 | 13.2 |
| Adipose | 1% | 32.2 | 19.4 |
| Adipose | 2% | 44.3 | 21.4 |
| Adipose | 5% | 50.8 | 22.1 |
| Adipose | 10% | 54.3 | 22.2 |
| Ileum | 1% | 23.2 | 15.5 |
| Ileum | 2% | 32.5 | 17.2 |
| Ileum | 5% | 42.0 | 18.0 |
| Ileum | 10% | 46.9 | 18.2 |
| **Mean** | 1% | 25.2 | 15.4 |
| **Mean** | 2% | 35.5 | 17.1 |
| **Mean** | 5% | 43.3 | 17.7 |
| **Mean** | 10% | 47.3 | 17.8 |

#### (D) FDR50 Within-Tissue

| Tissue | Target RDR | Peptide Increase (%) | Protein Increase (%) |
|---|---|---|---|
| Artery | 1% | 42.7 | 23.8 |
| Artery | 2% | 53.2 | 24.3 |
| Artery | 5% | 63.0 | 24.6 |
| Artery | 10% | 68.0 | 24.8 |
| Heart | 1% | 37.3 | 11.1 |
| Heart | 2% | 39.3 | 11.2 |
| Heart | 5% | 42.4 | 11.5 |
| Heart | 10% | 44.3 | 11.6 |
| Colon | 1% | 27.8 | 16.9 |
| Colon | 2% | 34.2 | 17.2 |
| Colon | 5% | 41.5 | 17.5 |
| Colon | 10% | 45.9 | 17.5 |
| Liver | 1% | 23.9 | 12.8 |
| Liver | 2% | 27.4 | 13.0 |
| Liver | 5% | 32.1 | 13.1 |
| Liver | 10% | 34.7 | 13.3 |
| Adipose | 1% | 38.9 | 20.9 |
| Adipose | 2% | 48.2 | 21.9 |
| Adipose | 5% | 54.0 | 22.2 |
| Adipose | 10% | 57.1 | 22.5 |
| Ileum | 1% | 30.9 | 17.0 |
| Ileum | 2% | 37.0 | 17.7 |
| Ileum | 5% | 44.0 | 18.0 |
| Ileum | 10% | 48.4 | 18.2 |
| **Mean** | 1% | 33.6 | 17.1 |
| **Mean** | 2% | 39.9 | 17.5 |
| **Mean** | 5% | 46.2 | 17.8 |
| **Mean** | 10% | 49.7 | 18.0 |

Percentage increase across 6 tissues compared to DIA-NN baseline (1% FDR, fast-gradient search). Peptide increase was calculated as (additional peptides / baseline peptides) × 100 for each sample. Protein group with at least one rescued peptide increase was calculated from DIA-NN Protein.Group annotations as (new protein groups / baseline protein groups) × 100 for each sample. Per-tissue values represent the mean across all samples within each tissue. Rows correspond to target RDR levels (1%, 2%, 5%, and 10%). **(A, C) Cross-tissue:** Models trained on all samples from the 5 other tissues, then applied to the held-out test tissue. **(B, D) Within-tissue:** Models trained on $n-1$ samples from the same tissue, then applied to the remaining held-out sample. **FDR20/FDR50:** FDR selected on DIA-NN (20% or 50%) used to recover additional peptides; both training and inference datasets contain peptides identified at this threshold.

**Table S4. Statistical comparison of within-tissue versus cross-tissue performance.** Number of additional peptides recovered by PeptiDIA per held-out sample (mean ± SD) for within-tissue and cross-tissue models, at each target reference-discordant rate (RDR: 1%, 2%, 5%, 10%). Δ(W−C) is the mean difference between strategies (within-tissue minus cross-tissue; green = within-tissue higher, red = cross-tissue higher). p-values are from two-tailed paired t-tests comparing within- versus cross-tissue recovery, paired by held-out sample; these correspond to the significance markers shown in Figures 3 and 4. (A) FDR20 models (trained on the DIA-NN 20% FDR candidate pool); (B) FDR50 models (DIA-NN 50% FDR candidate pool).

**A**

| Tissue | Target RDR | Within-tissue (mean ± SD) | Cross-tissue (mean ± SD) | Δ (W−C) | p (paired t) | Sig. |
|---|---|---|---|---|---|---|
| **Artery** | 1% | 306 ± 91 | 342 ± 108 | -36 | 0.008 | ** |
| | 2% | 388 ± 118 | 397 ± 126 | -9 | 0.082 | ns |
| | 5% | 450 ± 141 | 445 ± 141 | +5 | 0.065 | ns |
| | 10% | 475 ± 151 | 467 ± 147 | +8 | 2.5e-04 | *** |
| **Heart** | 1% | 517 ± 56 | 366 ± 49 | +151 | 1.3e-05 | *** |
| | 2% | 548 ± 59 | 444 ± 56 | +104 | 7.7e-06 | *** |
| | 5% | 580 ± 61 | 521 ± 56 | +59 | 2.4e-05 | *** |
| | 10% | 596 ± 62 | 558 ± 61 | +38 | 4.0e-06 | *** |
| **Colon** | 1% | 415 ± 109 | 398 ± 105 | +17 | 0.019 | * |
| | 2% | 491 ± 134 | 497 ± 128 | -6 | 0.202 | ns |
| | 5% | 586 ± 148 | 587 ± 149 | -0 | 0.814 | ns |
| | 10% | 628 ± 154 | 626 ± 153 | +2 | 0.225 | ns |
| **Liver** | 1% | 411 ± 128 | 342 ± 104 | +68 | 4.8e-04 | *** |
| | 2% | 461 ± 145 | 424 ± 138 | +37 | 2.0e-04 | *** |
| | 5% | 526 ± 170 | 501 ± 164 | +25 | 5.0e-04 | *** |
| | 10% | 551 ± 179 | 536 ± 176 | +15 | 1.2e-04 | *** |
| **Adipose** | 1% | 288 ± 22 | 274 ± 24 | +14 | 0.010 | ** |
| | 2% | 348 ± 23 | 331 ± 24 | +17 | 5.1e-05 | *** |
| | 5% | 377 ± 31 | 361 ± 27 | +16 | 6.7e-05 | *** |
| | 10% | 390 ± 32 | 380 ± 32 | +11 | 1.8e-05 | *** |
| **Ileum** | 1% | 528 ± 159 | 500 ± 147 | +28 | 0.004 | ** |
| | 2% | 619 ± 178 | 589 ± 171 | +30 | 9.3e-05 | *** |
| | 5% | 711 ± 193 | 694 ± 193 | +17 | 7.4e-05 | *** |
| | 10% | 755 ± 204 | 742 ± 202 | +13 | 3.8e-05 | *** |

**B**

| Tissue | Target RDR | Within-tissue (mean ± SD) | Cross-tissue (mean ± SD) | Δ (W−C) | p (paired t) | Sig. |
|---|---|---|---|---|---|---|
| **Artery** | 1% | 373 ± 118 | 368 ± 121 | +5 | 0.619 | ns |
| | 2% | 465 ± 149 | 464 ± 152 | +1 | 0.849 | ns |
| | 5% | 550 ± 174 | 536 ± 173 | +14 | 9.0e-05 | *** |
| | 10% | 594 ± 190 | 574 ± 184 | +20 | 1.3e-05 | *** |
| **Heart** | 1% | 642 ± 75 | 276 ± 40 | +366 | 2.6e-06 | *** |
| | 2% | 686 ± 81 | 477 ± 64 | +208 | 1.5e-05 | *** |
| | 5% | 740 ± 85 | 620 ± 80 | +121 | 3.3e-05 | *** |
| | 10% | 776 ± 87 | 677 ± 84 | +99 | 2.0e-05 | *** |
| **Colon** | 1% | 471 ± 141 | 361 ± 108 | +110 | 5.1e-06 | *** |
| | 2% | 580 ± 167 | 541 ± 153 | +39 | 4.7e-05 | *** |
| | 5% | 703 ± 188 | 676 ± 186 | +27 | 6.7e-06 | *** |
| | 10% | 779 ± 205 | 765 ± 204 | +14 | 2.2e-05 | *** |
| **Liver** | 1% | 472 ± 168 | 326 ± 106 | +146 | 0.001 | ** |
| | 2% | 543 ± 195 | 459 ± 159 | +83 | 0.002 | ** |
| | 5% | 635 ± 220 | 587 ± 209 | +48 | 4.2e-05 | *** |
| | 10% | 686 ± 231 | 646 ± 227 | +40 | 2.2e-05 | *** |
| **Adipose** | 1% | 320 ± 40 | 264 ± 31 | +56 | 3.0e-04 | *** |
| | 2% | 396 ± 38 | 364 ± 34 | +32 | 1.8e-05 | *** |
| | 5% | 443 ± 43 | 417 ± 41 | +26 | 1.1e-06 | *** |
| | 10% | 469 ± 52 | 446 ± 50 | +23 | 1.8e-07 | *** |
| **Ileum** | 1% | 594 ± 177 | 446 ± 136 | +147 | 2.3e-06 | *** |
| | 2% | 710 ± 207 | 624 ± 184 | +85 | 2.7e-06 | *** |
| | 5% | 845 ± 231 | 806 ± 228 | +39 | 5.4e-07 | *** |
| | 10% | 929 ± 246 | 901 ± 249 | +29 | 1.5e-06 | *** |

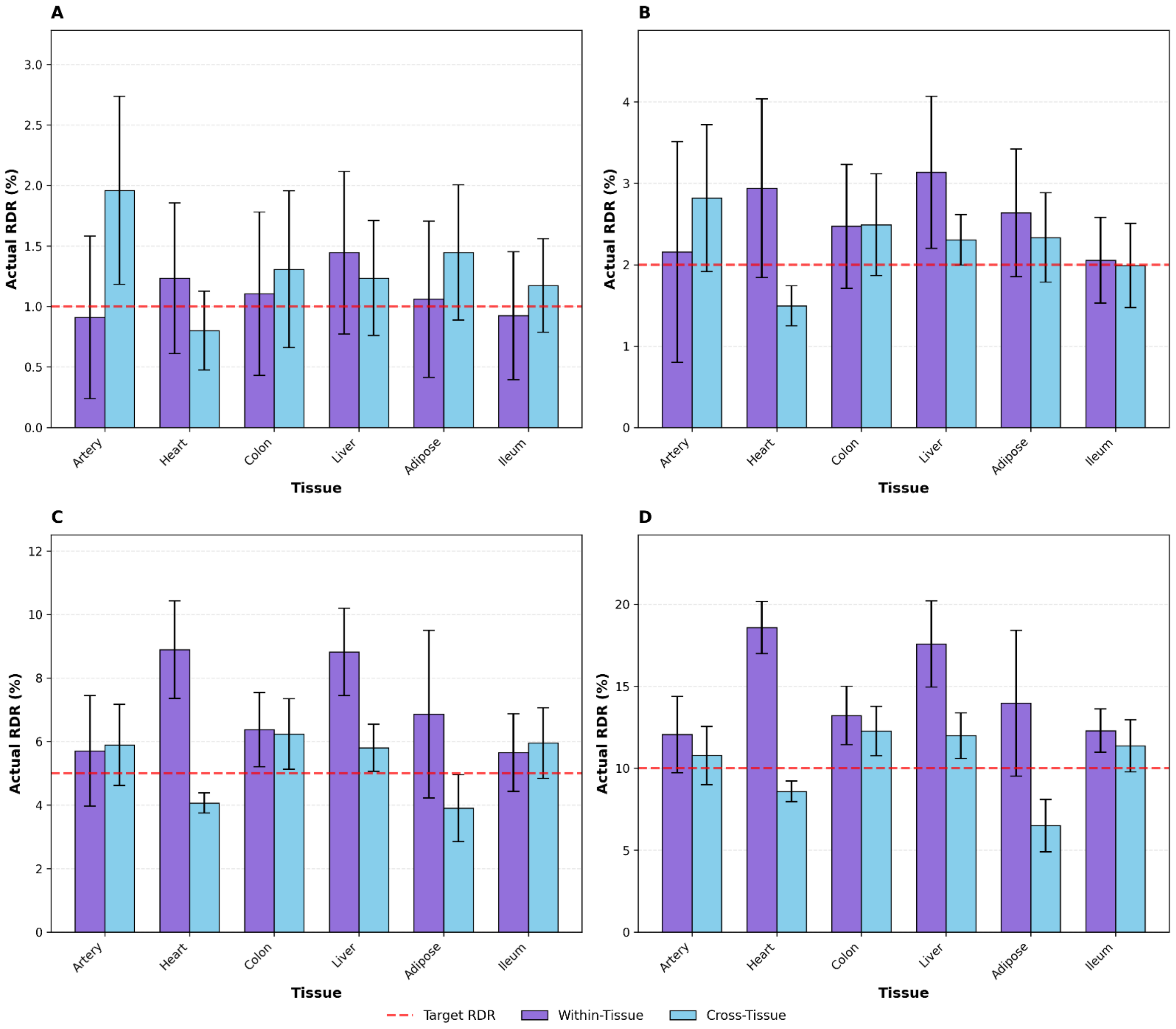


**Figure S3. FDR20 model: RDR calibration across target thresholds.** Bar plots showing the actual RDR achieved by PeptiDIA for each tissue at **(A)** 1%, **(B)** 2%, **(C)** 5%, and **(D)** 10% target RDR. Machine learning models were trained using additional peptides recovered by running DIA-NN at 20% FDR. Within-tissue models (purple) were trained on n-1 samples from the same tissue and tested on the held-out sample. Cross-tissue models (blue) were trained on all samples from the 5 other tissues and tested on the held-out tissue. Red dashed line indicates the target RDR. Error bars represent standard deviation. Values below the target line indicate conservative RDR control; values above indicate liberal RDR control.

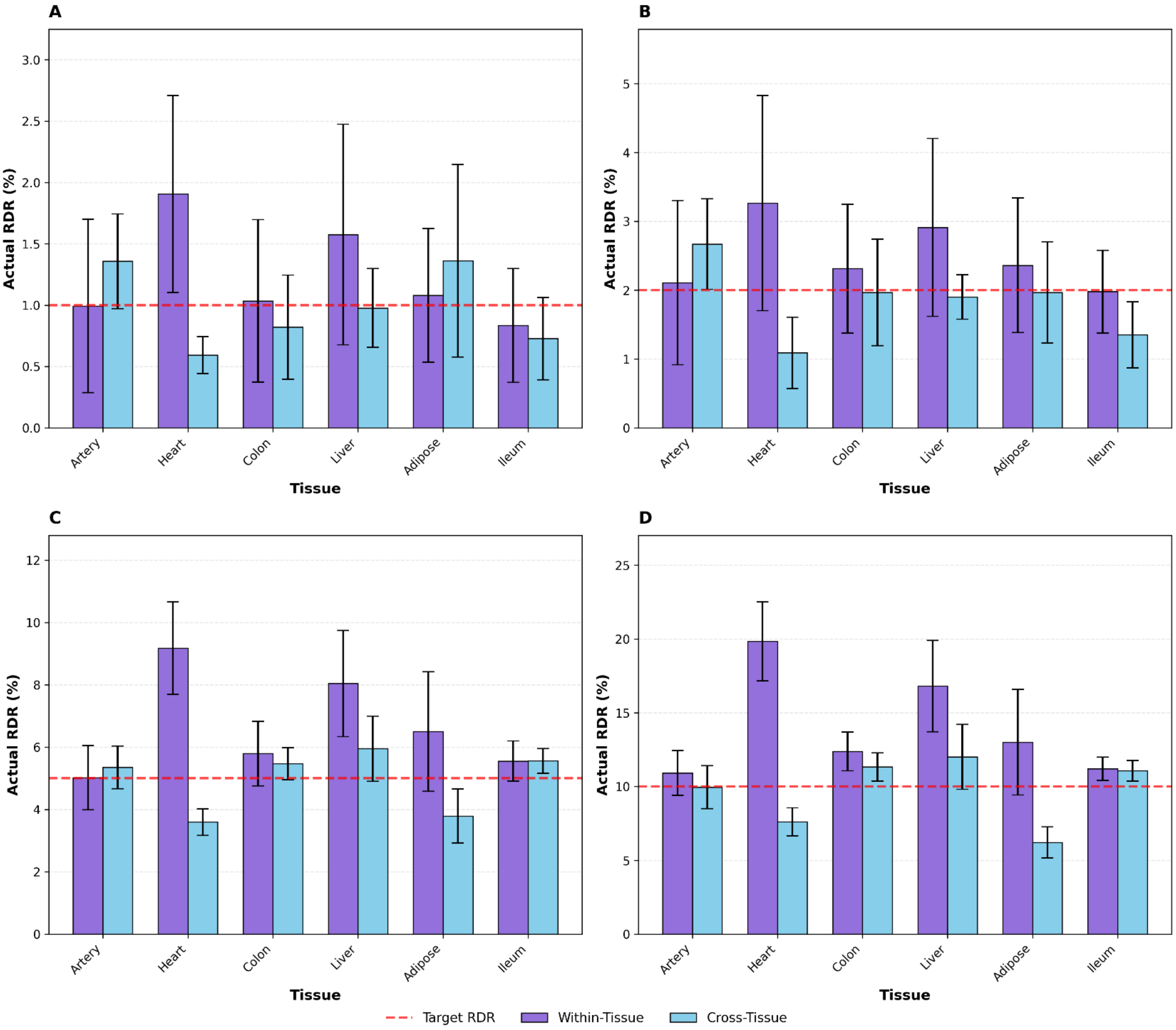


**Figure S4. FDR50 model: RDR calibration across target thresholds.** Bar plots showing the actual RDR achieved by PeptiDIA for each tissue at **(A)** 1%, **(B)** 2%, **(C)** 5%, and **(D)** 10% target RDR. Machine learning models were trained using additional peptides recovered by running DIA-NN at 50% FDR. Within-tissue models (purple) were trained on n-1 samples from the same tissue and tested on the held-out sample. Cross-tissue models (blue) were trained on all samples from the 5 other tissues and tested on the held-out tissue. Red dashed line indicates the target RDR. Error bars represent standard deviation. Values below the target line indicate conservative RDR control; values above indicate liberal RDR control.

### Table S5. PeptiDIA vs DIA-NN Comparison at Matched Peptide Counts

#### (A) FDR20 Cross-Tissue

| Tissue | Target RDR | Add. Pep. | ML RDR (%) | DIA-NN FDR (%) |
|---|---|---|---|---|
| Artery | 1% | 342 | 2.0 | 17.8 |
| Artery | 2% | 397 | 2.8 | 19.8 |
| Artery | 5% | 445 | 5.9 | 21.8 |
| Artery | 10% | 467 | 10.8 | 22.5 |
| Heart | 1% | 366 | 0.8 | 11.9 |
| Heart | 2% | 444 | 1.5 | 14.9 |
| Heart | 5% | 521 | 4.1 | 17.5 |
| Heart | 10% | 558 | 8.6 | 18.6 |
| Colon | 1% | 398 | 1.3 | 17.1 |
| Colon | 2% | 497 | 2.5 | 20.7 |
| Colon | 5% | 587 | 6.2 | 24.2 |
| Colon | 10% | 626 | 12.3 | 25.6 |
| Liver | 1% | 342 | 1.2 | 19.0 |
| Liver | 2% | 424 | 2.3 | 21.1 |
| Liver | 5% | 501 | 5.8 | 24.0 |
| Liver | 10% | 536 | 12.0 | 25.3 |
| Adipose | 1% | 274 | 1.4 | 14.2 |
| Adipose | 2% | 331 | 2.3 | 17.7 |
| Adipose | 5% | 361 | 3.9 | 19.6 |
| Adipose | 10% | 380 | 6.5 | 20.7 |
| Ileum | 1% | 500 | 1.2 | 16.7 |
| Ileum | 2% | 589 | 2.0 | 19.3 |
| Ileum | 5% | 694 | 6.0 | 22.7 |
| Ileum | 10% | 742 | 11.4 | 24.3 |
| **Mean** | 1% | 371 | 1.3 | 16.1 |
| **Mean** | 2% | 447 | 2.2 | 18.9 |
| **Mean** | 5% | 518 | 5.3 | 21.6 |
| **Mean** | 10% | 551 | 10.3 | 22.8 |

#### (B) FDR20 Within-Tissue

| Tissue | Target RDR | Add. Pep. | ML RDR (%) | DIA-NN FDR (%) |
|---|---|---|---|---|
| Artery | 1% | 306 | 0.9 | 16.4 |
| Artery | 2% | 388 | 2.2 | 19.1 |
| Artery | 5% | 450 | 5.7 | 21.8 |
| Artery | 10% | 475 | 12.0 | 22.9 |
| Heart | 1% | 517 | 1.2 | 17.4 |
| Heart | 2% | 548 | 2.9 | 18.2 |
| Heart | 5% | 580 | 8.9 | 19.7 |
| Heart | 10% | 596 | 18.6 | 20.0 |
| Colon | 1% | 415 | 1.1 | 17.6 |
| Colon | 2% | 491 | 2.5 | 20.5 |
| Colon | 5% | 586 | 6.4 | 24.1 |
| Colon | 10% | 628 | 13.2 | 25.6 |
| Liver | 1% | 411 | 1.4 | 20.9 |
| Liver | 2% | 461 | 3.1 | 22.9 |
| Liver | 5% | 526 | 8.8 | 24.8 |
| Liver | 10% | 551 | 17.6 | 26.0 |
| Adipose | 1% | 288 | 1.1 | 14.9 |
| Adipose | 2% | 348 | 2.6 | 18.6 |
| Adipose | 5% | 377 | 6.9 | 20.6 |
| Adipose | 10% | 390 | 14.0 | 21.2 |
| Ileum | 1% | 528 | 0.9 | 17.5 |
| Ileum | 2% | 619 | 2.1 | 20.3 |
| Ileum | 5% | 711 | 5.7 | 23.2 |
| Ileum | 10% | 755 | 12.3 | 24.8 |
| **Mean** | 1% | 411 | 1.1 | 17.4 |
| **Mean** | 2% | 476 | 2.6 | 19.9 |
| **Mean** | 5% | 538 | 7.1 | 22.4 |
| **Mean** | 10% | 566 | 14.6 | 23.4 |

#### (C) FDR50 Cross-Tissue

| Tissue | Target RDR | Add. Pep. | ML RDR (%) | DIA-NN FDR (%) |
|---|---|---|---|---|
| Artery | 1% | 368 | 1.4 | 18.5 |
| Artery | 2% | 464 | 2.7 | 22.3 |
| Artery | 5% | 536 | 5.3 | 25.6 |
| Artery | 10% | 574 | 10.0 | 27.8 |
| Heart | 1% | 276 | 0.6 | 8.9 |
| Heart | 2% | 477 | 1.1 | 15.9 |
| Heart | 5% | 620 | 3.6 | 21.2 |
| Heart | 10% | 677 | 7.6 | 23.4 |
| Colon | 1% | 361 | 0.8 | 16.1 |
| Colon | 2% | 541 | 2.0 | 22.2 |
| Colon | 5% | 676 | 5.5 | 27.2 |
| Colon | 10% | 765 | 11.3 | 30.5 |
| Liver | 1% | 326 | 1.0 | 18.1 |
| Liver | 2% | 459 | 1.9 | 22.4 |
| Liver | 5% | 587 | 6.0 | 28.0 |
| Liver | 10% | 646 | 12.0 | 31.2 |
| Adipose | 1% | 264 | 1.4 | 13.8 |
| Adipose | 2% | 364 | 2.0 | 19.5 |
| Adipose | 5% | 417 | 3.8 | 22.8 |
| Adipose | 10% | 446 | 6.2 | 24.8 |
| Ileum | 1% | 446 | 0.7 | 15.5 |
| Ileum | 2% | 624 | 1.4 | 20.1 |
| Ileum | 5% | 806 | 5.6 | 26.3 |
| Ileum | 10% | 901 | 11.1 | 29.2 |
| **Mean** | 1% | 340 | 1.0 | 15.2 |
| **Mean** | 2% | 488 | 1.9 | 20.4 |
| **Mean** | 5% | 607 | 5.0 | 25.2 |
| **Mean** | 10% | 668 | 9.7 | 27.8 |

#### (D) FDR50 Within-Tissue

| Tissue | Target RDR | Add. Pep. | ML RDR (%) | DIA-NN FDR (%) |
|---|---|---|---|---|
| Artery | 1% | 373 | 1.0 | 18.5 |
| Artery | 2% | 465 | 2.1 | 22.3 |
| Artery | 5% | 550 | 5.0 | 26.5 |
| Artery | 10% | 594 | 10.9 | 28.9 |
| Heart | 1% | 642 | 1.2 | 22.1 |
| Heart | 2% | 686 | 2.6 | 23.9 |
| Heart | 5% | 740 | 7.8 | 26.3 |
| Heart | 10% | 776 | 16.4 | 27.7 |
| Colon | 1% | 471 | 1.0 | 19.6 |
| Colon | 2% | 580 | 2.3 | 23.6 |
| Colon | 5% | 703 | 5.8 | 28.0 |
| Colon | 10% | 779 | 12.4 | 31.1 |
| Liver | 1% | 472 | 1.6 | 23.1 |
| Liver | 2% | 543 | 2.9 | 25.7 |
| Liver | 5% | 635 | 8.0 | 30.6 |
| Liver | 10% | 686 | 16.8 | 32.8 |
| Adipose | 1% | 320 | 1.1 | 16.9 |
| Adipose | 2% | 396 | 2.4 | 21.3 |
| Adipose | 5% | 443 | 6.5 | 24.6 |
| Adipose | 10% | 469 | 13.0 | 26.5 |
| Ileum | 1% | 594 | 0.8 | 19.3 |
| Ileum | 2% | 710 | 2.0 | 23.0 |
| Ileum | 5% | 845 | 5.6 | 27.4 |
| Ileum | 10% | 929 | 11.2 | 30.1 |
| **Mean** | 1% | 479 | 1.1 | 19.9 |
| **Mean** | 2% | 563 | 2.4 | 23.3 |
| **Mean** | 5% | 653 | 6.5 | 27.2 |
| **Mean** | 10% | 706 | 13.5 | 29.5 |

Comparison of PeptiDIA ML RDR versus DIA-NN FDR when relaxing the Q-value threshold to recover the same number of additional peptides (Add. Pep.). **(A, C) Cross-tissue:** Models trained on all samples from the 5 other tissues, then applied to the held-out test tissue. **(B, D) Within-tissue:** Models trained on n-1 samples from the same tissue, then applied to the remaining held-out sample. **FDR20/FDR50:** Refers to the DIA-NN Q-value threshold (20% or 50%) used to recover additional peptides for training the machine learning models.

**Table S6. Quantification validation: Spearman correlations between ML-rescued peptides and ground truth**

**(A) FDR20 Within-Tissue**

| Tissue | Baseline | 1% RDR | 2% RDR | 5% RDR | 10% RDR |
|---|---|---|---|---|---|
| Artery | 0.853 | 0.622 | 0.594 | 0.482 | 0.341 |
| Heart | 0.866 | 0.620 | 0.543 | 0.265 | 0.172 |
| Colon | 0.822 | 0.626 | 0.568 | 0.566 | 0.370 |
| Liver | 0.845 | 0.617 | 0.571 | 0.440 | 0.398 |
| Adipose | 0.874 | 0.738 | 0.640 | 0.558 | 0.521 |
| Ileum | 0.816 | 0.614 | 0.564 | 0.486 | 0.337 |
| Mean | 0.846 | 0.640 | 0.580 | 0.466 | 0.357 |

**(B) FDR20 Cross-Tissue**

| Tissue | Baseline | 1% RDR | 2% RDR | 5% RDR | 10% RDR |
|---|---|---|---|---|---|
| Artery | 0.853 | 0.625 | 0.493 | 0.486 | 0.356 |
| Heart | 0.866 | 0.665 | 0.629 | 0.508 | 0.519 |
| Colon | 0.822 | 0.625 | 0.585 | 0.524 | 0.371 |
| Liver | 0.845 | 0.650 | 0.535 | 0.502 | 0.241 |
| Adipose | 0.874 | 0.745 | 0.696 | 0.602 | 0.394 |
| Ileum | 0.816 | 0.606 | 0.663 | 0.561 | 0.381 |
| Mean | 0.846 | 0.653 | 0.600 | 0.531 | 0.377 |

**(C) FDR50 Within-Tissue**

| Tissue | Baseline | 1% RDR | 2% RDR | 5% RDR | 10% RDR |
|---|---|---|---|---|---|
| Artery | 0.853 | 0.581 | 0.564 | 0.340 | 0.232 |
| Heart | 0.866 | 0.606 | 0.436 | -0.086 | -0.038 |
| Colon | 0.822 | 0.614 | 0.472 | 0.429 | 0.210 |
| Liver | 0.845 | 0.615 | 0.511 | 0.414 | 0.435 |
| Adipose | 0.874 | 0.742 | 0.600 | 0.516 | 0.404 |
| Ileum | 0.816 | 0.595 | 0.498 | 0.351 | 0.231 |
| Mean | 0.846 | 0.626 | 0.514 | 0.327 | 0.246 |

**(D) FDR50 Cross-Tissue**

| Tissue | Baseline | 1% RDR | 2% RDR | 5% RDR | 10% RDR |
|---|---|---|---|---|---|
| Artery | 0.853 | 0.638 | 0.500 | 0.335 | 0.197 |
| Heart | 0.866 | 0.671 | 0.662 | 0.545 | 0.560 |
| Colon | 0.822 | 0.630 | 0.584 | 0.388 | 0.336 |
| Liver | 0.845 | 0.653 | 0.573 | 0.337 | 0.032 |
| Adipose | 0.874 | 0.751 | 0.692 | 0.520 | 0.373 |
| Ileum | 0.816 | 0.614 | 0.568 | 0.463 | 0.190 |
| Mean | 0.846 | 0.660 | 0.597 | 0.431 | 0.281 |

Spearman correlation coefficients ($\rho$) between $\log_2$-transformed precursor intensities (Precursor.Quantity in DIA-NN) of peptides identified in fast-gradient (300SPD) and matched ground truth intensities from long-gradient (30SPD) runs. **Baseline:** High-confidence peptides identified at 1% DIA-NN FDR in the fast-gradient search. **1–10% RDR:** ML-rescued peptides at different target RDR thresholds. **(A, B)** Models trained using peptides recovered at 20% DIA-NN FDR. **(C, D)** Models trained using peptides recovered at 50% DIA-NN FDR. **Within-tissue:** Models trained on n-1 samples from the same tissue and tested on the held-out sample. **Cross-tissue:** Models trained on all samples from 5 other tissues and tested on the held-out tissue. Higher correlations indicate better quantitative accuracy.

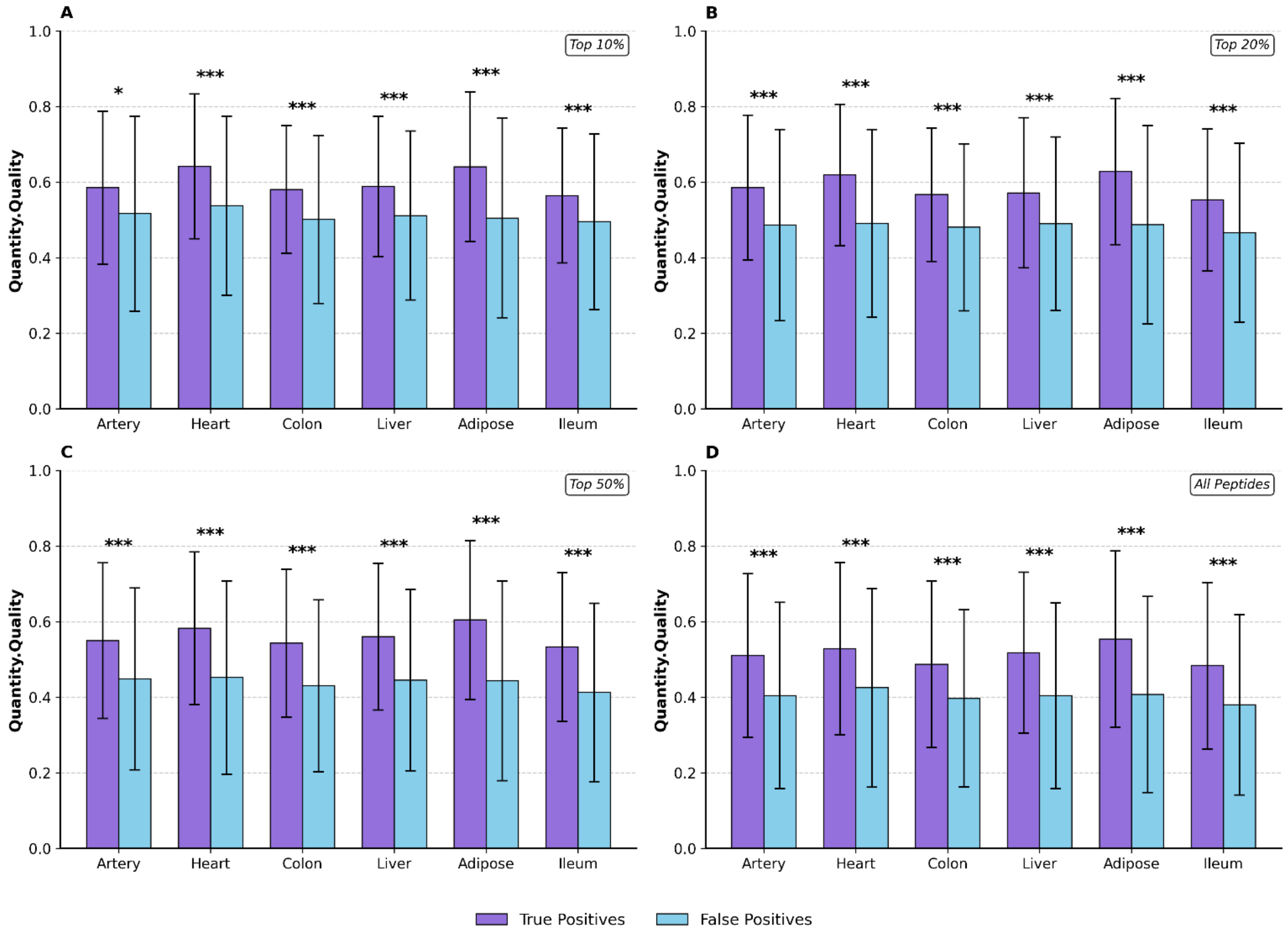


**Figure S5. DIA-NN Quantity Quality scores of True Positive versus False Positive Peptides (FDR20 training data).** Bar plots showing mean Quantity.Quality scores (± standard deviation) for True Positive peptides (purple; validated by long gradient ground truth) and False Positive peptides (cyan; not found in ground truth) across all tissues. Panels show results for peptides stratified by Q-value confidence: **(A) Top 10%**, **(B) Top 20%**, **(C) Top 50%**, and **(D) All peptides**. True Positives consistently exhibit significantly higher quality scores than False Positives across all tissues and FDR thresholds (Mann-Whitney U test; *$p < 0.05$, **$p < 0.01$, ***$p < 0.001$, ns = not significant).

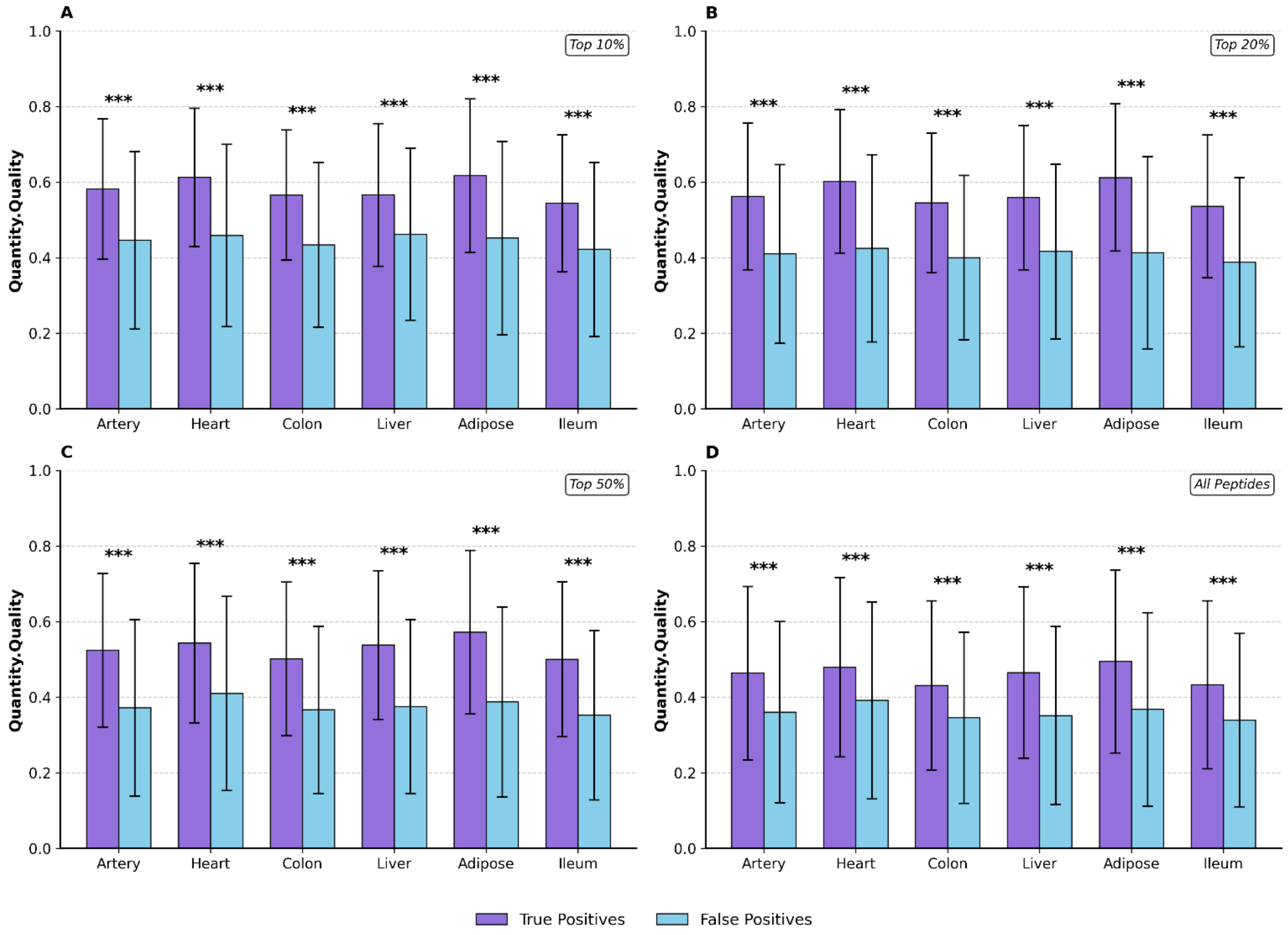


**Figure S6. DIA-NN Quantity Quality scores of True Positive versus False Positive Peptides (FDR50 training data).** Bar plots showing mean Quantity.Quality scores (± standard deviation) for True Positive peptides (purple; validated by long gradient ground truth) and False Positive peptides (cyan; not found in ground truth) across all tissues. Panels show results for peptides stratified by Q-value confidence: **(A) Top 10%**, **(B) Top 20%**, **(C) Top 50%**, and **(D) All peptides**. True Positives consistently exhibit significantly higher quality scores than False Positives across all tissues and FDR thresholds (Mann-Whitney U test; *$p < 0.05$, **$p < 0.01$, ***$p < 0.001$, ns = not significant).

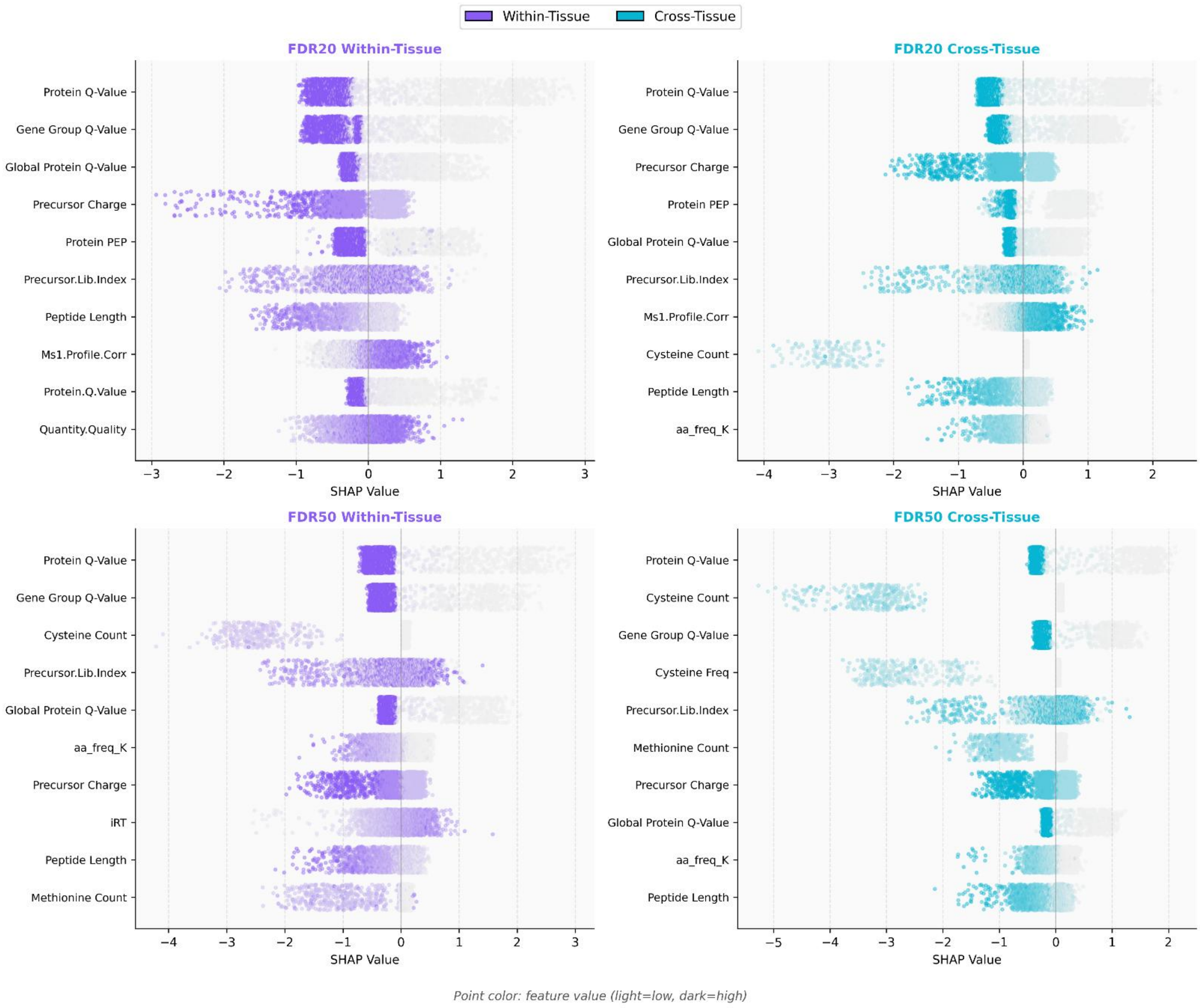


**Figure S7. SHAP (SHapley Additive exPlanations) analysis of feature contributions to XGBoost model predictions.** Beeswarm plots showing SHAP values for the top features across all four model conditions: FDR20 Within-Tissue (top-left), FDR20 Cross-Tissue (top-right), FDR50 Within-Tissue (bottom-left), and FDR50 Cross-Tissue (bottom-right). Each point represents a single peptide, with position along the x-axis indicating the magnitude and direction of the feature's impact on model prediction (positive SHAP values push predictions toward true positive classification; negative values toward false positive). Point color indicates the feature value (light = low, dark = high). Features are ranked by mean absolute SHAP value.

**Table S7. Complete feature importance rankings for XGBoost peptide validation models.** Feature importance values (gain) for all 87 features used in XGBoost gradient-boosted decision tree models across four validation conditions: FDR20 Within-Tissue, FDR20 Cross-Tissue, FDR50 Within-Tissue, and FDR50 Cross-Tissue. Gain represents the average reduction in training loss contributed by each feature across all tree splits. Values shown are mean ± standard deviation across all samples. Features are ranked by mean importance within each condition

| No. | Feature | Mean Importance | Std Importance | Condition |
|---|---|---|---|---|
| 1 | Global.PG.Q.Value | 167.38 | 40.26 | FDR20 Within-Tissue |
| 2 | PG.PEP | 86.73 | 21.1 | FDR20 Within-Tissue |
| 3 | PG.Q.Value | 77.3 | 28.57 | FDR20 Within-Tissue |
| 4 | GG.Q.Value | 59.18 | 39.35 | FDR20 Within-Tissue |
| 5 | aa_count_C | 19.61 | 7.23 | FDR20 Within-Tissue |
| 6 | aa_freq_C | 18.54 | 5.25 | FDR20 Within-Tissue |
| 7 | aa_count_M | 11.0 | 7.72 | FDR20 Within-Tissue |
| 8 | aa_freq_M | 8.19 | 4.63 | FDR20 Within-Tissue |
| 9 | Precursor.Charge | 6.14 | 1.26 | FDR20 Within-Tissue |
| 10 | sequence_length | 3.6 | 0.92 | FDR20 Within-Tissue |
| 11 | Global.Peptidoform.Q.Value | 3.29 | 1.21 | FDR20 Within-Tissue |
| 12 | aa_count_W | 3.25 | 1.07 | FDR20 Within-Tissue |
| 13 | Proteotypic | 3.08 | 1.3 | FDR20 Within-Tissue |
| 14 | Genes.MaxLFQ | 3.06 | 1.57 | FDR20 Within-Tissue |
| 15 | aa_count_R | 3.06 | 0.84 | FDR20 Within-Tissue |
| 16 | PG.MaxLFQ | 2.96 | 0.63 | FDR20 Within-Tissue |
| 17 | aa_count_H | 2.95 | 0.6 | FDR20 Within-Tissue |
| 18 | aa_freq_W | 2.83 | 0.88 | FDR20 Within-Tissue |
| 19 | Genes.MaxLFQ.Quality | 2.72 | 1.33 | FDR20 Within-Tissue |
| 20 | PEP | 2.67 | 1.02 | FDR20 Within-Tissue |
| 21 | aa_count_Y | 2.65 | 0.99 | FDR20 Within-Tissue |
| 22 | aa_freq_H | 2.61 | 0.31 | FDR20 Within-Tissue |
| 23 | zscore_Ms1.Area | 2.5 | 0.55 | FDR20 Within-Tissue |
| 24 | PG.MaxLFQ.Quality | 2.46 | 0.69 | FDR20 Within-Tissue |
| 25 | iRT | 2.43 | 0.37 | FDR20 Within-Tissue |
| 26 | aa_count_K | 2.39 | 0.25 | FDR20 Within-Tissue |
| 27 | Q.Value | 2.38 | 0.49 | FDR20 Within-Tissue |
| 28 | Ms1.Profile.Corr | 2.35 | 0.35 | FDR20 Within-Tissue |
| 29 | aa_freq_K | 2.24 | 0.35 | FDR20 Within-Tissue |
| 30 | log_Precursor.Quantity | 2.2 | 0.36 | FDR20 Within-Tissue |
| 31 | Ms1.Area | 2.12 | 0.4 | FDR20 Within-Tissue |
| 32 | aa_count_N | 2.05 | 0.31 | FDR20 Within-Tissue |
| 33 | RT.Start | 2.05 | 0.37 | FDR20 Within-Tissue |

| | | | | |
|---|---|---|---|---|
| 34 | Global.Q.Value | 2.04 | 0.43 | FDR20 Within-Tissue |
| 35 | aa_count_I | 1.99 | 0.43 | FDR20 Within-Tissue |
| 36 | aa_freq_P | 1.99 | 0.4 | FDR20 Within-Tissue |
| 37 | Predicted.RT | 1.96 | 0.49 | FDR20 Within-Tissue |
| 38 | aa_freq_Y | 1.95 | 0.29 | FDR20 Within-Tissue |
| 39 | Precursor.Lib.Index | 1.93 | 0.4 | FDR20 Within-Tissue |
| 40 | aa_count_E | 1.92 | 0.45 | FDR20 Within-Tissue |
| 41 | RT | 1.92 | 0.33 | FDR20 Within-Tissue |
| 42 | aa_freq_N | 1.92 | 0.23 | FDR20 Within-Tissue |
| 43 | aa_freq_F | 1.92 | 0.23 | FDR20 Within-Tissue |
| 44 | Genes.MaxLFQ.Unique | 1.88 | 0.32 | FDR20 Within-Tissue |
| 45 | Quantity.Quality | 1.85 | 0.38 | FDR20 Within-Tissue |
| 46 | Ms1.Apex.Area | 1.84 | 0.38 | FDR20 Within-Tissue |
| 47 | aa_count_S | 1.84 | 0.39 | FDR20 Within-Tissue |
| 48 | aa_count_F | 1.83 | 0.46 | FDR20 Within-Tissue |
| 49 | Genes.MaxLFQ.Unique.Quality | 1.82 | 0.41 | FDR20 Within-Tissue |
| 50 | aa_freq_I | 1.81 | 0.13 | FDR20 Within-Tissue |
| 51 | Precursor.Quantity | 1.8 | 0.13 | FDR20 Within-Tissue |
| 52 | aa_freq_Q | 1.79 | 0.34 | FDR20 Within-Tissue |
| 53 | Predicted.iRT | 1.79 | 0.35 | FDR20 Within-Tissue |
| 54 | aa_count_A | 1.79 | 0.45 | FDR20 Within-Tissue |
| 55 | aa_freq_D | 1.77 | 0.19 | FDR20 Within-Tissue |
| 56 | Precursor.Mz | 1.76 | 0.37 | FDR20 Within-Tissue |
| 57 | iIM | 1.76 | 0.31 | FDR20 Within-Tissue |
| 58 | Protein.Q.Value | 1.76 | 0.3 | FDR20 Within-Tissue |
| 59 | aa_count_L | 1.74 | 0.35 | FDR20 Within-Tissue |
| 60 | aa_count_P | 1.74 | 0.41 | FDR20 Within-Tissue |
| 61 | aa_count_T | 1.72 | 0.19 | FDR20 Within-Tissue |
| 62 | aa_freq_V | 1.72 | 0.3 | FDR20 Within-Tissue |
| 63 | RT.Stop | 1.71 | 0.35 | FDR20 Within-Tissue |
| 64 | aa_freq_R | 1.71 | 0.23 | FDR20 Within-Tissue |
| 65 | aa_freq_L | 1.67 | 0.47 | FDR20 Within-Tissue |
| 66 | aa_freq_G | 1.64 | 0.24 | FDR20 Within-Tissue |
| 67 | aa_freq_E | 1.64 | 0.27 | FDR20 Within-Tissue |
| 68 | aa_count_D | 1.62 | 0.6 | FDR20 Within-Tissue |
| 69 | Mass.Evidence | 1.59 | 0.4 | FDR20 Within-Tissue |
| 70 | aa_count_V | 1.58 | 0.22 | FDR20 Within-Tissue |
| 71 | Ms1.Normalised | 1.57 | 0.2 | FDR20 Within-Tissue |
| 72 | aa_freq_T | 1.57 | 0.23 | FDR20 Within-Tissue |
| 73 | aa_count_G | 1.56 | 0.28 | FDR20 Within-Tissue |

| | | | | |
|---|---|---|---|---|
| 74 | log_Ms1.Area | 1.55 | 0.32 | FDR20 Within-Tissue |
| 75 | aa_freq_S | 1.54 | 0.34 | FDR20 Within-Tissue |
| 76 | Channel.Evidence | 1.54 | 0.21 | FDR20 Within-Tissue |
| 77 | Precursor.Normalised | 1.54 | 0.24 | FDR20 Within-Tissue |
| 78 | aa_freq_A | 1.49 | 0.29 | FDR20 Within-Tissue |
| 79 | aa_count_Q | 1.47 | 0.3 | FDR20 Within-Tissue |
| 80 | FWHM | 1.4 | 0.18 | FDR20 Within-Tissue |
| 81 | Evidence | 1.4 | 0.12 | FDR20 Within-Tissue |
| 82 | Best.Fr.Mz | 1.33 | 0.14 | FDR20 Within-Tissue |
| 83 | Ms1.Total.Signal.After | 1.31 | 0.12 | FDR20 Within-Tissue |
| 84 | Ms1.Apex.Mz.Delta | 1.3 | 0.23 | FDR20 Within-Tissue |
| 85 | zscore_Precursor.Quantity | 1.25 | 0.37 | FDR20 Within-Tissue |
| 86 | Best.Fr.Mz.Delta | 1.21 | 0.27 | FDR20 Within-Tissue |
| 87 | Ms1.Total.Signal.Before | 1.18 | 0.14 | FDR20 Within-Tissue |
| 1 | Global.PG.Q.Value | 277.79 | 48.18 | FDR20 Cross-Tissue |
| 2 | PG.Q.Value | 197.4 | 20.05 | FDR20 Cross-Tissue |
| 3 | PG.PEP | 138.5 | 42.26 | FDR20 Cross-Tissue |
| 4 | GG.Q.Value | 136.93 | 44.64 | FDR20 Cross-Tissue |
| 5 | aa_count_C | 57.74 | 10.54 | FDR20 Cross-Tissue |
| 6 | aa_freq_C | 26.6 | 6.1 | FDR20 Cross-Tissue |
| 7 | aa_count_M | 25.52 | 6.67 | FDR20 Cross-Tissue |
| 8 | Precursor.Charge | 21.12 | 0.9 | FDR20 Cross-Tissue |
| 9 | aa_freq_M | 13.06 | 2.9 | FDR20 Cross-Tissue |
| 10 | Proteotypic | 9.25 | 0.76 | FDR20 Cross-Tissue |
| 11 | sequence_length | 8.24 | 0.75 | FDR20 Cross-Tissue |
| 12 | Global.Peptidoform.Q.Value | 8.16 | 0.92 | FDR20 Cross-Tissue |
| 13 | aa_count_W | 7.72 | 1.19 | FDR20 Cross-Tissue |
| 14 | aa_count_R | 7.71 | 0.63 | FDR20 Cross-Tissue |
| 15 | aa_count_H | 7.7 | 1.26 | FDR20 Cross-Tissue |
| 16 | PEP | 6.79 | 0.61 | FDR20 Cross-Tissue |
| 17 | aa_count_K | 6.73 | 0.63 | FDR20 Cross-Tissue |
| 18 | aa_count_Y | 6.68 | 1.37 | FDR20 Cross-Tissue |
| 19 | aa_freq_W | 6.63 | 0.58 | FDR20 Cross-Tissue |
| 20 | Ms1.Profile.Corr | 6.1 | 0.5 | FDR20 Cross-Tissue |
| 21 | aa_freq_H | 5.9 | 0.34 | FDR20 Cross-Tissue |
| 22 | aa_freq_K | 4.99 | 0.34 | FDR20 Cross-Tissue |
| 23 | log_Precursor.Quantity | 4.98 | 0.37 | FDR20 Cross-Tissue |
| 24 | iRT | 4.96 | 0.37 | FDR20 Cross-Tissue |
| 25 | Q.Value | 4.87 | 0.44 | FDR20 Cross-Tissue |
| 26 | PG.MaxLFQ.Quality | 4.67 | 0.42 | FDR20 Cross-Tissue |

| | | | | |
|---|---|---|---|---|
| 27 | Predicted.RT | 4.49 | 0.23 | FDR20 Cross-Tissue |
| 28 | aa_freq_P | 4.49 | 0.22 | FDR20 Cross-Tissue |
| 29 | aa_count_N | 4.43 | 0.8 | FDR20 Cross-Tissue |
| 30 | RT.Start | 4.42 | 0.32 | FDR20 Cross-Tissue |
| 31 | aa_freq_Y | 4.32 | 0.33 | FDR20 Cross-Tissue |
| 32 | Precursor.Lib.Index | 4.23 | 0.29 | FDR20 Cross-Tissue |
| 33 | zscore_Ms1.Area | 4.22 | 0.67 | FDR20 Cross-Tissue |
| 34 | aa_count_I | 4.16 | 0.83 | FDR20 Cross-Tissue |
| 35 | PG.MaxLFQ | 4.11 | 0.21 | FDR20 Cross-Tissue |
| 36 | aa_count_F | 3.99 | 0.52 | FDR20 Cross-Tissue |
| 37 | aa_count_P | 3.99 | 0.38 | FDR20 Cross-Tissue |
| 38 | Genes.MaxLFQ.Quality | 3.98 | 0.55 | FDR20 Cross-Tissue |
| 39 | Ms1.Apex.Area | 3.89 | 0.26 | FDR20 Cross-Tissue |
| 40 | Genes.MaxLFQ | 3.88 | 0.36 | FDR20 Cross-Tissue |
| 41 | Precursor.Quantity | 3.88 | 0.21 | FDR20 Cross-Tissue |
| 42 | Ms1.Area | 3.87 | 0.23 | FDR20 Cross-Tissue |
| 43 | Quantity.Quality | 3.82 | 0.26 | FDR20 Cross-Tissue |
| 44 | aa_freq_I | 3.82 | 0.12 | FDR20 Cross-Tissue |
| 45 | Precursor.Mz | 3.82 | 0.21 | FDR20 Cross-Tissue |
| 46 | aa_freq_N | 3.81 | 0.35 | FDR20 Cross-Tissue |
| 47 | aa_freq_R | 3.81 | 0.24 | FDR20 Cross-Tissue |
| 48 | aa_freq_F | 3.8 | 0.32 | FDR20 Cross-Tissue |
| 49 | Predicted.iRT | 3.77 | 0.24 | FDR20 Cross-Tissue |
| 50 | aa_freq_D | 3.73 | 0.24 | FDR20 Cross-Tissue |
| 51 | aa_count_S | 3.72 | 0.38 | FDR20 Cross-Tissue |
| 52 | iIM | 3.68 | 0.22 | FDR20 Cross-Tissue |
| 53 | RT | 3.66 | 0.18 | FDR20 Cross-Tissue |
| 54 | aa_freq_L | 3.61 | 0.26 | FDR20 Cross-Tissue |
| 55 | aa_count_D | 3.6 | 0.4 | FDR20 Cross-Tissue |
| 56 | aa_freq_V | 3.57 | 0.28 | FDR20 Cross-Tissue |
| 57 | aa_count_E | 3.53 | 0.22 | FDR20 Cross-Tissue |
| 58 | aa_count_T | 3.52 | 0.39 | FDR20 Cross-Tissue |
| 59 | aa_count_A | 3.51 | 0.28 | FDR20 Cross-Tissue |
| 60 | aa_freq_S | 3.5 | 0.31 | FDR20 Cross-Tissue |
| 61 | Global.Q.Value | 3.49 | 0.29 | FDR20 Cross-Tissue |
| 62 | aa_count_L | 3.47 | 0.27 | FDR20 Cross-Tissue |
| 63 | aa_freq_G | 3.47 | 0.16 | FDR20 Cross-Tissue |
| 64 | RT.Stop | 3.46 | 0.26 | FDR20 Cross-Tissue |
| 65 | aa_freq_Q | 3.45 | 0.25 | FDR20 Cross-Tissue |
| 66 | Protein.Q.Value | 3.45 | 0.19 | FDR20 Cross-Tissue |

| | | | | |
|---|---|---|---|---|
| 67 | aa_freq_A | 3.45 | 0.24 | FDR20 Cross-Tissue |
| 68 | Mass.Evidence | 3.43 | 0.24 | FDR20 Cross-Tissue |
| 69 | Genes.MaxLFQ.Unique | 3.41 | 0.19 | FDR20 Cross-Tissue |
| 70 | aa_count_G | 3.37 | 0.31 | FDR20 Cross-Tissue |
| 71 | aa_freq_T | 3.36 | 0.21 | FDR20 Cross-Tissue |
| 72 | aa_freq_E | 3.35 | 0.19 | FDR20 Cross-Tissue |
| 73 | aa_count_V | 3.24 | 0.37 | FDR20 Cross-Tissue |
| 74 | aa_count_Q | 3.21 | 0.18 | FDR20 Cross-Tissue |
| 75 | Ms1.Normalised | 3.12 | 0.19 | FDR20 Cross-Tissue |
| 76 | log_Ms1.Area | 3.09 | 0.18 | FDR20 Cross-Tissue |
| 77 | Precursor.Normalised | 3.09 | 0.24 | FDR20 Cross-Tissue |
| 78 | Genes.MaxLFQ.Unique.Quality | 2.99 | 0.22 | FDR20 Cross-Tissue |
| 79 | Channel.Evidence | 2.92 | 0.2 | FDR20 Cross-Tissue |
| 80 | zscore_Precursor.Quantity | 2.89 | 0.58 | FDR20 Cross-Tissue |
| 81 | Evidence | 2.78 | 0.12 | FDR20 Cross-Tissue |
| 82 | Ms1.Total.Signal.Before | 2.65 | 0.12 | FDR20 Cross-Tissue |
| 83 | FWHM | 2.64 | 0.1 | FDR20 Cross-Tissue |
| 84 | Ms1.Total.Signal.After | 2.62 | 0.04 | FDR20 Cross-Tissue |
| 85 | Best.Fr.Mz | 2.52 | 0.09 | FDR20 Cross-Tissue |
| 86 | Ms1.Apex.Mz.Delta | 2.52 | 0.12 | FDR20 Cross-Tissue |
| 87 | Best.Fr.Mz.Delta | 2.37 | 0.06 | FDR20 Cross-Tissue |
| 1 | Global.PG.Q.Value | 399.54 | 133.03 | FDR50 Within-Tissue |
| 2 | GG.Q.Value | 182.03 | 140.97 | FDR50 Within-Tissue |
| 3 | PG.Q.Value | 147.12 | 37.48 | FDR50 Within-Tissue |
| 4 | PG.PEP | 96.59 | 69.58 | FDR50 Within-Tissue |
| 5 | aa_count_C | 31.27 | 13.0 | FDR50 Within-Tissue |
| 6 | aa_freq_C | 28.09 | 11.33 | FDR50 Within-Tissue |
| 7 | aa_count_M | 17.52 | 11.42 | FDR50 Within-Tissue |
| 8 | aa_freq_M | 11.47 | 7.05 | FDR50 Within-Tissue |
| 9 | Global.Peptidoform.Q.Value | 9.5 | 5.5 | FDR50 Within-Tissue |
| 10 | Precursor.Charge | 9.0 | 2.68 | FDR50 Within-Tissue |
| 11 | PEP | 6.56 | 4.86 | FDR50 Within-Tissue |
| 12 | aa_count_R | 6.32 | 2.89 | FDR50 Within-Tissue |
| 13 | aa_count_K | 5.22 | 1.49 | FDR50 Within-Tissue |
| 14 | Proteotypic | 5.19 | 1.62 | FDR50 Within-Tissue |
| 15 | Q.Value | 4.99 | 1.72 | FDR50 Within-Tissue |
| 16 | aa_count_W | 4.85 | 1.48 | FDR50 Within-Tissue |
| 17 | sequence_length | 4.04 | 1.28 | FDR50 Within-Tissue |
| 18 | Global.Q.Value | 4.02 | 1.41 | FDR50 Within-Tissue |
| 19 | aa_freq_W | 3.48 | 1.09 | FDR50 Within-Tissue |

| | | | | |
|---|---|---|---|---|
| 20 | PG.MaxLFQ | 3.35 | 0.43 | FDR50 Within-Tissue |
| 21 | aa_count_H | 3.31 | 0.57 | FDR50 Within-Tissue |
| 22 | aa_freq_H | 3.16 | 0.57 | FDR50 Within-Tissue |
| 23 | Genes.MaxLFQ | 3.1 | 0.9 | FDR50 Within-Tissue |
| 24 | aa_freq_K | 3.03 | 0.55 | FDR50 Within-Tissue |
| 25 | iRT | 2.99 | 0.69 | FDR50 Within-Tissue |
| 26 | PG.MaxLFQ.Quality | 2.98 | 1.39 | FDR50 Within-Tissue |
| 27 | aa_count_Y | 2.97 | 0.8 | FDR50 Within-Tissue |
| 28 | Ms1.Profile.Corr | 2.77 | 0.36 | FDR50 Within-Tissue |
| 29 | log_Precursor.Quantity | 2.71 | 0.35 | FDR50 Within-Tissue |
| 30 | Precursor.Lib.Index | 2.57 | 0.62 | FDR50 Within-Tissue |
| 31 | aa_freq_P | 2.56 | 0.62 | FDR50 Within-Tissue |
| 32 | Genes.MaxLFQ.Quality | 2.54 | 0.74 | FDR50 Within-Tissue |
| 33 | Genes.MaxLFQ.Unique | 2.45 | 0.21 | FDR50 Within-Tissue |
| 34 | zscore_Ms1.Area | 2.44 | 0.45 | FDR50 Within-Tissue |
| 35 | aa_count_N | 2.43 | 0.79 | FDR50 Within-Tissue |
| 36 | aa_count_F | 2.43 | 0.41 | FDR50 Within-Tissue |
| 37 | aa_freq_Y | 2.43 | 0.49 | FDR50 Within-Tissue |
| 38 | Predicted.RT | 2.42 | 0.59 | FDR50 Within-Tissue |
| 39 | aa_freq_I | 2.38 | 0.27 | FDR50 Within-Tissue |
| 40 | aa_freq_R | 2.37 | 0.49 | FDR50 Within-Tissue |
| 41 | RT.Start | 2.36 | 0.7 | FDR50 Within-Tissue |
| 42 | aa_freq_F | 2.32 | 0.43 | FDR50 Within-Tissue |
| 43 | aa_freq_N | 2.28 | 0.45 | FDR50 Within-Tissue |
| 44 | aa_count_L | 2.26 | 0.36 | FDR50 Within-Tissue |
| 45 | Ms1.Area | 2.26 | 0.27 | FDR50 Within-Tissue |
| 46 | aa_count_S | 2.22 | 0.44 | FDR50 Within-Tissue |
| 47 | aa_freq_D | 2.21 | 0.45 | FDR50 Within-Tissue |
| 48 | Precursor.Mz | 2.2 | 0.42 | FDR50 Within-Tissue |
| 49 | Quantity.Quality | 2.19 | 0.52 | FDR50 Within-Tissue |
| 50 | aa_freq_V | 2.18 | 0.44 | FDR50 Within-Tissue |
| 51 | Mass.Evidence | 2.16 | 0.46 | FDR50 Within-Tissue |
| 52 | aa_count_E | 2.16 | 0.29 | FDR50 Within-Tissue |
| 53 | aa_freq_Q | 2.16 | 0.54 | FDR50 Within-Tissue |
| 54 | aa_count_V | 2.15 | 0.17 | FDR50 Within-Tissue |
| 55 | Predicted.iRT | 2.14 | 0.45 | FDR50 Within-Tissue |
| 56 | aa_count_A | 2.14 | 0.58 | FDR50 Within-Tissue |
| 57 | RT | 2.13 | 0.3 | FDR50 Within-Tissue |
| 58 | aa_count_I | 2.12 | 0.44 | FDR50 Within-Tissue |
| 59 | aa_count_T | 2.12 | 0.37 | FDR50 Within-Tissue |

| | | | | |
|---|---|---|---|---|
| 60 | aa_freq_L | 2.11 | 0.6 | FDR50 Within-Tissue |
| 61 | iIM | 2.11 | 0.32 | FDR50 Within-Tissue |
| 62 | aa_freq_G | 2.11 | 0.22 | FDR50 Within-Tissue |
| 63 | aa_count_P | 2.1 | 0.45 | FDR50 Within-Tissue |
| 64 | Ms1.Apex.Area | 2.1 | 0.52 | FDR50 Within-Tissue |
| 65 | aa_freq_S | 2.08 | 0.41 | FDR50 Within-Tissue |
| 66 | Genes.MaxLFQ.Unique.Quality | 2.05 | 0.5 | FDR50 Within-Tissue |
| 67 | aa_freq_E | 2.03 | 0.35 | FDR50 Within-Tissue |
| 68 | Precursor.Quantity | 2.03 | 0.35 | FDR50 Within-Tissue |
| 69 | aa_count_G | 2.01 | 0.42 | FDR50 Within-Tissue |
| 70 | RT.Stop | 2.01 | 0.32 | FDR50 Within-Tissue |
| 71 | Protein.Q.Value | 2.0 | 0.3 | FDR50 Within-Tissue |
| 72 | aa_freq_T | 2.0 | 0.32 | FDR50 Within-Tissue |
| 73 | log_Ms1.Area | 1.98 | 0.29 | FDR50 Within-Tissue |
| 74 | Ms1.Normalised | 1.97 | 0.18 | FDR50 Within-Tissue |
| 75 | aa_freq_A | 1.97 | 0.37 | FDR50 Within-Tissue |
| 76 | aa_count_Q | 1.96 | 0.44 | FDR50 Within-Tissue |
| 77 | aa_count_D | 1.96 | 0.56 | FDR50 Within-Tissue |
| 78 | Precursor.Normalised | 1.89 | 0.21 | FDR50 Within-Tissue |
| 79 | Channel.Evidence | 1.88 | 0.23 | FDR50 Within-Tissue |
| 80 | FWHM | 1.72 | 0.22 | FDR50 Within-Tissue |
| 81 | Evidence | 1.72 | 0.23 | FDR50 Within-Tissue |
| 82 | Best.Fr.Mz | 1.69 | 0.23 | FDR50 Within-Tissue |
| 83 | Ms1.Apex.Mz.Delta | 1.67 | 0.27 | FDR50 Within-Tissue |
| 84 | Ms1.Total.Signal.After | 1.6 | 0.12 | FDR50 Within-Tissue |
| 85 | zscore_Precursor.Quantity | 1.58 | 0.34 | FDR50 Within-Tissue |
| 86 | Best.Fr.Mz.Delta | 1.55 | 0.28 | FDR50 Within-Tissue |
| 87 | Ms1.Total.Signal.Before | 1.5 | 0.21 | FDR50 Within-Tissue |
| 1 | Global.PG.Q.Value | 822.0 | 184.24 | FDR50 Cross-Tissue |
| 2 | PG.Q.Value | 527.49 | 106.36 | FDR50 Cross-Tissue |
| 3 | GG.Q.Value | 238.54 | 101.86 | FDR50 Cross-Tissue |
| 4 | PG.PEP | 183.71 | 76.06 | FDR50 Cross-Tissue |
| 5 | aa_count_C | 114.56 | 14.93 | FDR50 Cross-Tissue |
| 6 | aa_freq_C | 56.75 | 8.64 | FDR50 Cross-Tissue |
| 7 | aa_count_M | 47.67 | 12.38 | FDR50 Cross-Tissue |
| 8 | Precursor.Charge | 30.82 | 2.24 | FDR50 Cross-Tissue |
| 9 | Global.Peptidoform.Q.Value | 28.44 | 7.16 | FDR50 Cross-Tissue |
| 10 | aa_freq_M | 22.18 | 6.61 | FDR50 Cross-Tissue |
| 11 | aa_count_R | 16.98 | 1.7 | FDR50 Cross-Tissue |
| 12 | Q.Value | 16.1 | 1.85 | FDR50 Cross-Tissue |

| | | | | |
|---|---|---|---|---|
| 13 | Proteotypic | 15.53 | 1.2 | FDR50 Cross-Tissue |
| 14 | PEP | 14.91 | 3.36 | FDR50 Cross-Tissue |
| 15 | aa_count_K | 13.18 | 1.37 | FDR50 Cross-Tissue |
| 16 | Global.Q.Value | 10.8 | 1.14 | FDR50 Cross-Tissue |
| 17 | aa_count_W | 10.69 | 1.13 | FDR50 Cross-Tissue |
| 18 | sequence_length | 10.0 | 0.99 | FDR50 Cross-Tissue |
| 19 | aa_count_H | 9.32 | 1.57 | FDR50 Cross-Tissue |
| 20 | aa_freq_W | 8.12 | 0.46 | FDR50 Cross-Tissue |
| 21 | Ms1.Profile.Corr | 7.41 | 0.32 | FDR50 Cross-Tissue |
| 22 | aa_freq_H | 7.4 | 0.34 | FDR50 Cross-Tissue |
| 23 | aa_freq_K | 7.19 | 0.52 | FDR50 Cross-Tissue |
| 24 | aa_count_Y | 6.9 | 0.92 | FDR50 Cross-Tissue |
| 25 | iRT | 6.9 | 0.59 | FDR50 Cross-Tissue |
| 26 | Precursor.Lib.Index | 6.34 | 0.45 | FDR50 Cross-Tissue |
| 27 | RT.Start | 6.02 | 0.45 | FDR50 Cross-Tissue |
| 28 | Predicted.RT | 5.98 | 0.44 | FDR50 Cross-Tissue |
| 29 | aa_freq_P | 5.93 | 0.41 | FDR50 Cross-Tissue |
| 30 | aa_freq_R | 5.75 | 0.26 | FDR50 Cross-Tissue |
| 31 | aa_freq_Y | 5.64 | 0.38 | FDR50 Cross-Tissue |
| 32 | PG.MaxLFQ.Quality | 5.59 | 0.4 | FDR50 Cross-Tissue |
| 33 | PG.MaxLFQ | 5.58 | 0.34 | FDR50 Cross-Tissue |
| 34 | log_Precursor.Quantity | 5.56 | 0.51 | FDR50 Cross-Tissue |
| 35 | aa_count_N | 5.52 | 0.55 | FDR50 Cross-Tissue |
| 36 | Quantity.Quality | 5.35 | 0.4 | FDR50 Cross-Tissue |
| 37 | aa_freq_I | 5.24 | 0.26 | FDR50 Cross-Tissue |
| 38 | aa_count_P | 5.23 | 0.31 | FDR50 Cross-Tissue |
| 39 | aa_freq_N | 5.22 | 0.48 | FDR50 Cross-Tissue |
| 40 | zscore_Ms1.Area | 5.18 | 0.36 | FDR50 Cross-Tissue |
| 41 | Precursor.Mz | 5.17 | 0.33 | FDR50 Cross-Tissue |
| 42 | RT | 5.16 | 0.31 | FDR50 Cross-Tissue |
| 43 | Predicted.iRT | 5.14 | 0.38 | FDR50 Cross-Tissue |
| 44 | Mass.Evidence | 5.14 | 0.33 | FDR50 Cross-Tissue |
| 45 | Ms1.Area | 5.11 | 0.15 | FDR50 Cross-Tissue |
| 46 | iIM | 5.1 | 0.31 | FDR50 Cross-Tissue |
| 47 | Genes.MaxLFQ | 5.09 | 0.17 | FDR50 Cross-Tissue |
| 48 | aa_freq_F | 5.08 | 0.29 | FDR50 Cross-Tissue |
| 49 | aa_freq_S | 5.06 | 0.34 | FDR50 Cross-Tissue |
| 50 | aa_count_S | 5.05 | 0.34 | FDR50 Cross-Tissue |
| 51 | Genes.MaxLFQ.Quality | 5.03 | 0.92 | FDR50 Cross-Tissue |
| 52 | aa_freq_L | 5.02 | 0.41 | FDR50 Cross-Tissue |

| | | | | |
|---|---|---|---|---|
| 53 | aa_freq_V | 4.89 | 0.33 | FDR50 Cross-Tissue |
| 54 | aa_count_F | 4.87 | 0.31 | FDR50 Cross-Tissue |
| 55 | aa_freq_G | 4.87 | 0.24 | FDR50 Cross-Tissue |
| 56 | aa_freq_D | 4.86 | 0.33 | FDR50 Cross-Tissue |
| 57 | aa_count_I | 4.81 | 0.36 | FDR50 Cross-Tissue |
| 58 | Ms1.Apex.Area | 4.78 | 0.32 | FDR50 Cross-Tissue |
| 59 | aa_count_A | 4.74 | 0.45 | FDR50 Cross-Tissue |
| 60 | aa_freq_E | 4.73 | 0.35 | FDR50 Cross-Tissue |
| 61 | aa_freq_Q | 4.65 | 0.27 | FDR50 Cross-Tissue |
| 62 | aa_freq_A | 4.61 | 0.34 | FDR50 Cross-Tissue |
| 63 | aa_count_V | 4.61 | 0.38 | FDR50 Cross-Tissue |
| 64 | aa_count_D | 4.6 | 0.52 | FDR50 Cross-Tissue |
| 65 | aa_count_T | 4.59 | 0.26 | FDR50 Cross-Tissue |
| 66 | aa_count_G | 4.58 | 0.32 | FDR50 Cross-Tissue |
| 67 | aa_count_E | 4.56 | 0.46 | FDR50 Cross-Tissue |
| 68 | aa_count_Q | 4.51 | 0.43 | FDR50 Cross-Tissue |
| 69 | Protein.Q.Value | 4.49 | 0.28 | FDR50 Cross-Tissue |
| 70 | RT.Stop | 4.47 | 0.31 | FDR50 Cross-Tissue |
| 71 | aa_freq_T | 4.46 | 0.32 | FDR50 Cross-Tissue |
| 72 | aa_count_L | 4.46 | 0.27 | FDR50 Cross-Tissue |
| 73 | log_Ms1.Area | 4.42 | 0.36 | FDR50 Cross-Tissue |
| 74 | Ms1.Normalised | 4.36 | 0.23 | FDR50 Cross-Tissue |
| 75 | Precursor.Quantity | 4.35 | 0.33 | FDR50 Cross-Tissue |
| 76 | Genes.MaxLFQ.Unique | 4.32 | 0.21 | FDR50 Cross-Tissue |
| 77 | Precursor.Normalised | 4.25 | 0.27 | FDR50 Cross-Tissue |
| 78 | Genes.MaxLFQ.Unique.Quality | 4.03 | 0.27 | FDR50 Cross-Tissue |
| 79 | zscore_Precursor.Quantity | 3.99 | 0.39 | FDR50 Cross-Tissue |
| 80 | Channel.Evidence | 3.98 | 0.15 | FDR50 Cross-Tissue |
| 81 | Evidence | 3.82 | 0.12 | FDR50 Cross-Tissue |
| 82 | Ms1.Total.Signal.Before | 3.65 | 0.18 | FDR50 Cross-Tissue |
| 83 | Ms1.Apex.Mz.Delta | 3.62 | 0.16 | FDR50 Cross-Tissue |
| 84 | Ms1.Total.Signal.After | 3.61 | 0.26 | FDR50 Cross-Tissue |
| 85 | FWHM | 3.59 | 0.04 | FDR50 Cross-Tissue |
| 86 | Best.Fr.Mz | 3.46 | 0.1 | FDR50 Cross-Tissue |
| 87 | Best.Fr.Mz.Delta | 3.2 | 0.05 | FDR50 Cross-Tissue |

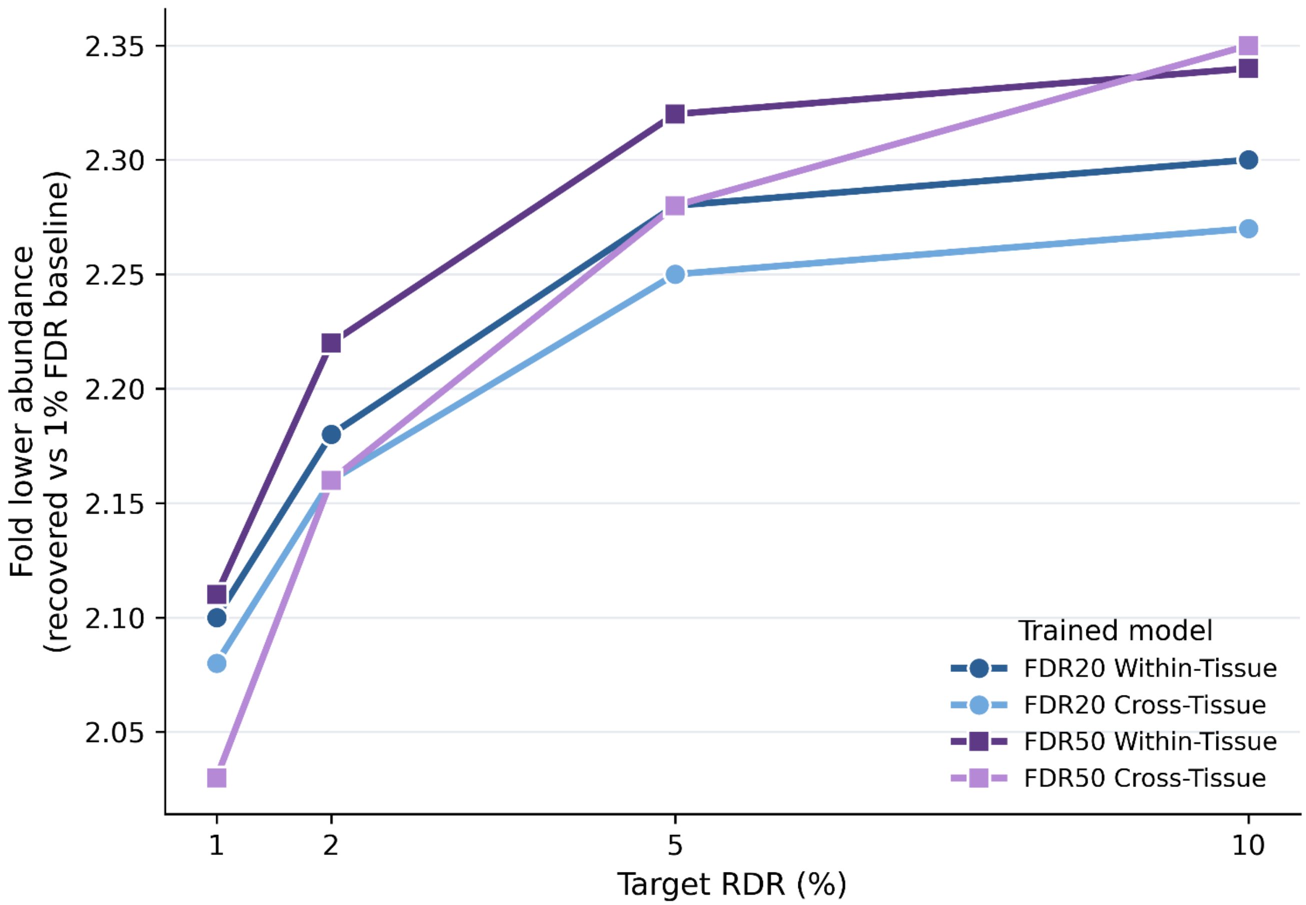


**Figure S8. Recovered peptides are of lower abundance than the 1% FDR baseline.** Median fold-lower abundance of PeptiDIA-recovered peptides relative to the 1% FDR baseline (DIA-NN) as a function of target reference-discordant rate (RDR), shown for all four model conditions: FDR20 Within-Tissue, FDR20 Cross-Tissue, FDR50 Within-Tissue, and FDR50 Cross-Tissue. Each series shows the mean fold difference in peptide abundance across tissues at each target RDR (1%, 2%, 5%, 10%); higher values indicate that newly recovered peptides are progressively lower in abundance.
